# From criticality to cognitive effort: scale-invariant EEG dynamics supporting cognitive flexibility are suppressed by effort

**DOI:** 10.64898/2026.05.10.724092

**Authors:** Li Xin Lim, Abhimanyu Bhardwaj, Arthur-Ervin Avramiea, Klaus Linkenkaer-Hansen, Asako Mitsuto, Lily Tran, Woodrow Shew, Andrew Westbrook

## Abstract

The critical brain hypothesis contends that brains operate near a phase transition where excitation and inhibition are balanced, enabling neural dynamics to rapidly adapt and reorganize for cognitive demands. Allocating control resources to maintain stable task representations likely shifts brains away from criticality. Here, we test whether proximity to criticality indexes the balance between flexible adaptation and effortful task engagement. To do so, we adapt a time-resolved measure of scale invariance in EEG amplitude fluctuations (*d2*), capable of quantifying distance-to-criticality under non-stationary conditions – as during cognitive tasks. We benchmark our measure using ground-truth simulations of a neural mass model and show that *d2* is lowest when excitation and inhibition are balanced. Next, we apply *d2* to data collected during a task-switching paradigm and find that more demanding trials increased deviation from criticality, whereas greater flexibility, faster responses, and higher accuracy occurred closer to criticality. These effects were region-specific: deviation at posterior electrodes predicted worse performance, while deviations at frontal midline electrodes predicted better performance. Together, these results suggest that deviations from criticality reflect both cognitive load and effort exertion, highlighting EEG amplitude scale-invariance as a sensitive marker of adaptive neural dynamics under cognitive demand.

## 1. Introduction

A key question in cognitive neuroscience is how the brain mediates the tradeoff between the engagement of cognitive resources needed to perform tasks while remaining flexibly adaptable to varying cognitive demands. Cognitive effort involves the processes by which the brain adjusts the level of engagement required to meet task demands relative to available processing capacity (Kahneman, 1973). Effort also acts as a regulatory system that allocates limited resources based on the expected value and the perceived costs of exerting control (Botvinick and Braver, 2015; Kool and Botvinick, 2018; Shenhav et al., 2017). Increasing cognitive load sensitizes information processing to task demands, such that even minor changes can alter how information is organized, integrated, and used to guide behavior (Collins et al., 2017; Crossley et al., 2018; Lim et al., 2025). This naturally raises the broader question of what neural conditions enable the brain to support such efficient and adaptive information processing. This question becomes especially important when considering large-scale neural dynamics that reflect the brain’s delicate balance between stability and flexibility under limited cognitive resources.

A key aspect of this balance is the brain’s capacity to process and integrate information across different spatial and temporal scales, allowing adaptive responses to changing environments. Brain functions cover a wide range of timescales, from quick perceptual-motor events to longer-lasting higher-order cognitive processes over seconds. The critical brain hypothesis (Beggs, 2008; Hengen & Shew, 2025) suggests that healthy neural activity operates near a critical point transition between ordered and disordered phases. This naturally produces fluctuations that are invariant across these timescales, and the range of this scale-invariance widens as the system approaches criticality (Jones et al., 2023). At a critical point, emergent dynamics naturally arising from the interplay of excitatory and inhibitory activity create a balanced regime where activity neither dampens too quickly, as in subcritical systems, nor grows uncontrollably, as in supercritical systems. This balance enables the network to maintain the broadest dynamic range, support optimal information processing and transmission, and increase sensitivity to external stimuli, thereby promoting flexible and efficient computation (Ezaki et al., 2020; Gautam et al., 2015; Müller et al., 2025; Shew et al., 2011; Shriki & Yellin, 2016). Evidence of criticality appears across neural scales, from in vitro and in vivo neuronal activity (Dahmen et al., 2019; Gireesh & Plenz, 2008; Hahn et al., 2017; Ma et al., 2019) to large-scale recordings such as EEG, MEG, and fMRI, characterized by features like power-law avalanches, long-range correlations, and scale-invariant spectral patterns (Arviv et al., 2015; A. E. Avramiea et al., 2022; Kitzbichler et al., 2009; Linkenkaer-Hansen et al., 2001).

Importantly, distance-to-criticality is not fixed; it reorganizes following perturbations and is influenced by pharmacological interventions (Fekete et al., 2018; Maschke et al., 2024) or sleep deprivation (Colombo et al., 2016; Meisel et al., 2017). Thus, when a task imposes greater demands requiring more selective engagement of neural resources, the brain may move away from criticality, deviating from its default, highly flexible state, to focus on the specific computations needed. Converging neuroimaging evidence supports this perspective. fMRI studies have shown that the fractal scaling of BOLD signals systematically decreases with increased cognitive load, including more demanding perceptual decisions, novel tasks, and higher working memory loads (Barnes et al., 2009; Churchill et al., 2016), suggesting that suppression of scale-invariance serves as a marker of cognitive demand. Importantly, it may also index effort exerted. EEG research has demonstrated monotonic reductions in scale-invariance, quantified by the Hurst exponent (which measures fractality), as working memory load increases, even beyond capacity limits, suggesting that effort, beyond merely load, suppresses scale-free activity (Kardan et al., 2020). Similarly, better sustained attention performance was linked to reduced long-range temporal correlations during tasks relative to rest, suggesting greater suppression of scale-free activity (Irrmischer et al., 2018). Another EEG study (Fagerholm et al., 2015) found that cortical cascade distributions follow near-critical power laws during rest but shift toward subcritical dynamics during task performance, with this deviation predicting both sensorimotor BOLD activation and faster reaction times. Together, these findings suggest that cognitive demand, and the exertion of cognitive effort both reliably push neural activity away from the flexible, high-dynamic-range regime of near-criticality toward states aligned with task demands.

While prior work suggests that cognitive demands influence proximity to criticality, testing this has been limited by the fact that empirical assays of criticality typically require long-duration, recordings, assuming stationarity, but periods of cognitive effort are often short in experiments. Thus, testing how cognitive effort affects proximity to criticality requires methods with higher temporal resolution. Recently, Sooter et al. (2025) introduced a method to measure proximity to criticality based on characteristics of autoregressive models fit to short time series. After fitting autoregressive models, distance-to-criticality (*d2*) is calculated as the Euclidean distance between the fitted model and the nearest scale-invariant autoregressive model. *d2* leverages temporal renormalization group (tRG) theory to identify autoregressive models that generate temporally scale-invariant fluctuations (i.e. tRG fixed points) expected at criticality. Because autoregressive models can be estimated reliably from short time series, the method permits time-resolved measurements of distance to criticality. This approach is essential for capturing within- and between-trial fluctuations in criticality, as these rapid changes occur on a timescales that are far too short for conventional, long-window metrics to detect. When applied to spike recordings in mouse visual cortex, *d2* revealed that while the relaxed awake state is closest to criticality, relative to heightened arousal and deep sleep, distance-to-criticality rapidly fluctuates with phasic bursts of body movement (Sooter et al., 2025).

While *d2* was first used to track distance-to-criticality in spiking neurons, the present study adapts the tRG-derived metric to track between- and within-trial dynamics in distance-to-criticality, based on amplitude fluctuations in EEG scalp recordings. Applying *d2* to amplitude fluctuations is supported by a recent study (Fontenele et al., 2025) in a computational model of critical oscillations, which shows that the critical oscillatory dynamics become evident after isolating the amplitude envelope. Our aim was to capture rapid shifts in brain criticality during a task-switching paradigm where participants must flexibly alternate among cognitive operations. We used *d2* to test the hypotheses that operating closer to criticality facilitates task flexibility (Hellyer et al., 2015; Simola et al., 2017), and enhances the brain’s ability to adapt and select appropriate cognitive strategies. In contrast, transient deviations from criticality may reflect the selective engagement of resources needed for demanding, effortful task execution. By applying *d2* in this paradigm, we aimed to test fine-grained, temporally-resolved predictions about how criticality supports adaptive cognitive effort.

## 2. Methodology

### 2.1 Participants

88 participants (44 female, 37 male, 7 gender not reported) were recruited from Brown University and surrounding communities. Participants were all healthy young adults (ages 18–43, mean = 22 years, SD = 5.0 years) without any psychiatric or neurological disorders, and not currently taking any psychiatric medications. Five participants with excessively noisy recordings were excluded leaving *N* = 83 for analyses. Each participant received monetary compensation for their participation. After providing informed consent, participants were prepared for EEG recordings and completed a set of computer-based tasks lasting approximately 3 hours in total. Research protocols were approved by Brown University’s Institutional Review Board (Protocol number: 2106003016) and were conducted in full compliance with institutional and ethical guidelines for research involving human participants, including the Declaration of Helsinki of 1975.

### 2.2 Task and Materials

#### 2.2.1 Task description

A task-switching paradigm (Armbruster et al., 2012; Jos et al., 2025), as shown in Fig. 1A, involving both Regular and non-Regular (forced-switch, distractor, and voluntary-switch trials) trials, was used to assess relationships between criticality, cognitive flexibility, and task demand. Each trial consisted of one or two numbers presented in relation to a central sine-wave grating (circular line pattern). Starting with an inter-trial interval, a Gabor patch was displayed at the center of the screen at the start of each trial. Participants performed one of the two tasks: the odd/even task (indicating whether a number was odd or even) or the high/low task (indicating whether a number was greater than or less than 5). Concurrent with the onset of the number stimulus, the Gabor orientation changed, cuing which task should be performed on that trial. In Regular trials (Gabor grating oriented at 30° from horizontal), only a single number appeared in a typical location (e.g., below the Gabor patch). In all other trial types, numbers were displayed both above and below the grating. On forced-switch trials (Gabor grating at 80°), participants were required to respond to the task and number in the atypical location, on distractor trials (Gabor grating at 30°), participants were required to ignore the number in the atypical location, and on voluntary-switch trials (Gabor grating at 50°) they selected the task they wanted to perform. Responses were made via keyboard keys ‘f’, ‘d’, ‘j’, and ‘k’. The two adjacent keys were assigned to the same task (either the high/low task or the odd/even task), and key assignments and task positions were counterbalanced across participants. Each trial allowed a maximum response window of 2.6 s followed by a uniformly-sampled jitter from 1.0–3.0 s, introducing temporal variability and reducing predictability.

**Figure 1.**
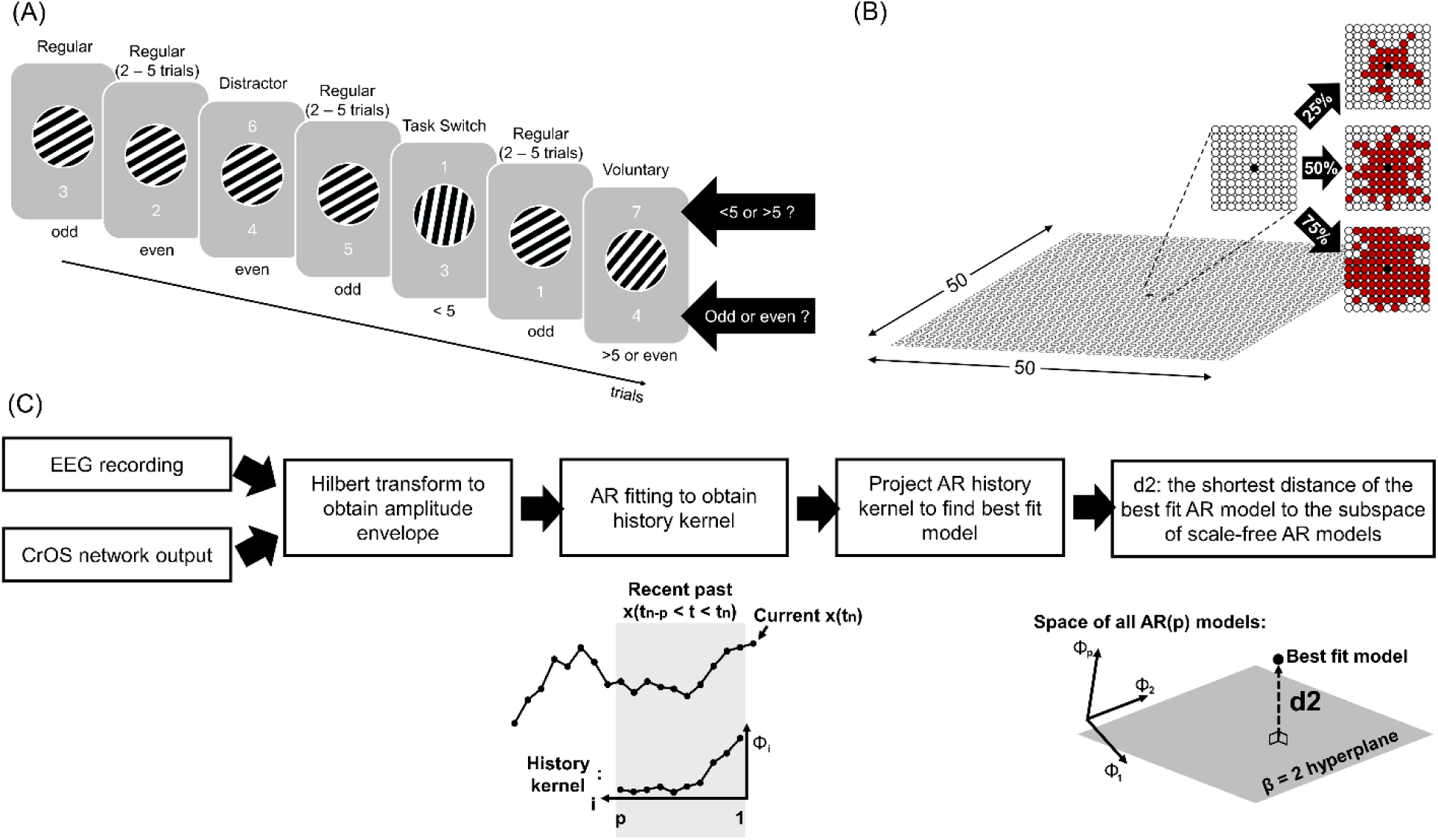
(A) Schematic of the task-switching paradigm. (B) Example simulations from the CrOS model (Poil et al., 2012) on a 50×50 grid with 25%, 50% and 75% excitatory connectivity. (C) Illustration of the *d2* computation procedure (Sooter et al., 2025), where either EEG recordings or CROS network outputs serve as input signals. The amplitude envelope of the input is extracted, an autoregressive (AR) model is fitted to the envelope to estimate the history kernel, and *d2* is defined as the Euclidean distance between an AR history kernel, and the nearest AR model produced by a scale-invariant system.

#### 2.2.2 CROS model simulation

For ground-truth simulations, we used the probabilistic firing-rate (CRitical OScillations, or CROS) neural mass model which undergoes a phase transition from an inhibition-dominant subcritical, to an excitation dominant supercritical phase, as a function of excitatory to inhibitory connectivity (A. E. Avramiea et al., 2022; A.-E. Avramiea et al., 2025; Poil et al., 2012). At criticality, the CROS model exhibits power-law distributed avalanches and long-range temporal correlations in oscillatory amplitudes. The network consisted of integrate-and-fire neurons arranged on a 50×50 open grid (shown in Fig. 1B), with 75% excitatory and 25% inhibitory neurons. Connectivity was defined by two parameters, CE and CI, representing the percentage of neurons within a local neighborhood that each excitatory and inhibitory neuron, respectively, are connected to. These parameters were systematically varied from 25% to 100% in 5% increments, and three independent network realizations were generated for each CE and CI combination. External stimuli were injected by a group of spike-generating neurons delivering periodic, jittered (delivered at intervals averaging 1s, jittered by ±250ms), excitatory input to a randomly-chosen subset of excitatory neurons, increasing the excitatory input current of the target neurons, which in turn propagated activity through the recurrent network. The number of stimulated neurons was determined by the parameter *stimulus_size* (e.g., with *stimulus_size* = 25, approximately 1% of all neurons, or ∼1.33% of excitatory neurons, were directly driven). These external spikes acted as brief excitatory perturbation, mimicking sensory or task-related inputs to the network.

### 2.3 Procedure

#### 2.3.1 Task description

The present experiment was carried out within the scope of a larger investigation into criticality and working memory. In the resting phase which preceded the task, participants maintained open eyes while fixating on a central fixation cross for five minutes. Participants then completed a training phase consisting of 30 trials per block, repeated as needed until 80% accuracy was achieved on the non-Regular trials to ensure adequate task comprehension. After training, participants proceeded to the main experiment, which consisted of 210 trials. Because the final practice block was repeated for participants who did not meet the accuracy criterion, some individuals contributed more than 210 trials to the analysis.

#### 2.3.2 CROS model simulation

For each network defined by a specific combination of CE and CI, the simulation lasted 10 minutes (with a sample size of 300,000 and a sampling frequency of 500 Hz). The spiking activity of all excitatory and inhibitory neurons was summed to obtain the network output. Notably, the simulated network output from the CROS model exhibits oscillatory dynamics, a feature commonly observed in neural time series recordings such as EEG.

### 2.4 Data Preprocessing and Analysis

#### 2.4.1 EEG preprocessing

Participants were seated upright in a sound-attenuated room for EEG recordings using a 64-channel cap with active electrodes and a BrainVision amplifier. 1024 Hz recordings were collected throughout the resting period and task, then down-sampled to 512 Hz. Additionally, data were high-pass filtered at 1Hz with a Hamming windowed sinc FIR filter (using the pop_eegfiltnew function in EEGLAB). Bad channels were automatically detected with clean_artifacts, which removed channels showing low correlation (<0.65) with neighboring channels or excessive line noise (>4 SD). This led to an average of 3.0 (SD = 2.5) channels being removed. These channels were then interpolated using pop_interp and re-referenced to the average with pop_reref. The data were further processed with clean_artifacts, using only the BurstCriterion parameter set to 30, and re-referenced again to the average. Next, independent components analysis (ICA) was performed with pop_runica to identify components. According to the ICLabel algorithm, components likely to be muscle, eye, heart, line noise, or channel noise artifacts with a probability greater than 0.7 were identified and rejected.

#### 2.4.2 Pupil dilation analysis

Pupil diameter was recorded binocularly using Eyelogic LogicOne 250, with a sampling rate of 250 Hz. For subsequent analyses, only the right-eye signal was used. A temporal window from 0.650 to 2.00 s after stimulus onset was selected to capture peak pupil dilation response on Regular trials, while 1.50 to 2.85 s post-stimulus was used for non-Regular trials. Within this window, the mean pupil diameter was computed for each trial, yielding a single value per trial for statistical analysis.

#### 2.4.3 Distance to criticality (*d2*)

Distance-to-criticality (Figure 1C) was estimated using the tRG-based scale-invariance adapted from the method introduced by (Sooter et al., 2025). We adapted this measure to evaluate scale-invariance in fluctuations of the amplitude envelope of EEG data or the CROS model’s population signal, computed by the Hilbert transform. We used the amplitude envelope rather than raw signal because raw signal contains oscillations which are scale-dependent, anti-correlated timeseries. In contrast, the amplitude envelope reflects coordinated activity across scales of neuronal assemblies.

To evaluate fluctuations in distance-to-criticality over time, we computed *d2* in 2.5 s sliding windows, based on a window size that maximizes sensitivity to fluctuations in non-stationary environments, while preserving stability in *d2* estimates in EEG data sampled at 512 Hz (see Supplemental Figure S1). For on-task data, windows were slid with a 98% overlap from -2.25 to 2.5 s relative to stimulus onset. For the resting-state data, we applied our sliding window over the entire EEG recording (5 min). For CROS model simulations, we applied sliding windows to the amplitude envelope of summed network output in each combination of CE and CI across the entire time series (10 minutes simulation of 500 Hz sampling frequency).

Each envelope segment (2.5 s, corresponding to 1280 samples at a sampling rate of 512 Hz) was divided into bins of 250 samples each. For each bin, an autoregression model with order 25 (AR(25)) was fit to the data after subtracting the within-bin mean to focus on temporal fluctuations around local activity levels. The best fitting AR(25) was computed using the mean-centered data, up to the model order (number of successive time lags). AR coefficients were fit using the Yule-Walker method, which finds the set of AR parameters whose implied autocorrelation structure best matches the empirical, mean-centered data. The resulting AR history kernel provided an estimate of temporal dependencies in the envelope fluctuations.

A complete mathematical description of the d2 analysis is reported elsewhere (Sooter et al., 2025), but we provide a brief account here. To evaluate the distance-to-criticality, we compared the empirical best-fit AR(25) history kernel against the set of kernels of all critical AR(25) models. A critical AR(25) model is defined as one that generates temporally scale-invariant fluctuations. Previously theory established that, in the 25-dimensional space of all AR(25) models, there is a 25×1 dimensional hyperplane of critical AR(25) models. Within that set of critical AR(25) models there are different types of criticality, defined by power spectra with 1/f^β^ power-law form, with exponents β = 2, 4, …, up to twice the model order. For each β, a kernel was constructed using binomial coefficients with alternating signs to capture the expected structure of critical dynamics. The empirical AR kernel was then projected onto these theoretical kernels, and the Euclidean distance between the empirical kernel and its closest critical projection was calculated. This Euclidean distance served as the distance-to-criticality measure (*d2*), with larger values of *d2* indicating greater divergence of amplitude fluctuations from critical temporal organization.

#### 2.4.4 Detrended fluctuation analysis (DFA)

DFA (Peng et al., 1994) was applied to both network output signals from CROS model simulations and resting-state EEG recorded during the experiment. For each combination of CE and CI from CROS model simulations, we extracted activity in the alpha frequency range (8.30 – 13.4 Hz) using band-pass filtering. Following (Poil et al., 2012) we first obtained the amplitude envelope of the filtered signals by Hilbert transform, then cumulatively summed the amplitude envelope. Next, we segmented the cumulative sum into logarithmically-spaced window sizes, each window was linearly detrended, and the scaling exponent was obtained from the slope of the log-log relation between window size and mean magnitude of the fluctuation computed in each detrended segment.

#### 2.4.5 Kappa

To quantify how closely avalanche-size distributions followed the expected critical power-law, the κ index (Shew et al., 2009) was calculated following the procedure in (Poil et al., 2012). κ is calculated as the mean difference between the cumulative distribution of avalanche sizes, derived from the amplitude envelope of the network output signal in CROS model simulations, and a reference power law with an exponent of -1.5. This difference is assessed at 50 logarithmically spaced points across the observed size range, followed by the addition of 1.0. Values of κ < 1 indicate subcritical dynamics, κ > 1 indicate supercritical dynamics, and κ ≈ 1 reflects near-critical organization.

#### 2.4.6 Time-lagged auto-mutual information

Mutual information was used to quantify temporal information propagation in the network signal. For each simulation, mutual information was computed between the signal at time *t* and the upcoming value *t*+*τ*, for lags τ = 1 to 25 (following the AR order for d2), using a 20-bin discretization (log2N + 1 = 19.19). Mutual information values were then averaged across lags to obtain a single summary measure of temporal information transfer for each simulation and subsequently averaged across runs.

## 3. Results

### 3.1 *d2* tracks excitation-inhibition balance in ground-truth simulations

To test the fidelity with which *d2* indexes criticality, we computed *d2* using simulations of a ground-truth model of coupled excitatory and inhibitory neurons: the CROS model (A.-E. Avramiea et al., 2025; Poil et al., 2012), which undergoes a phase transition. Evidence of criticality in this model includes the emergence of long-range temporal correlations in amplitude fluctuations and power-law distributed neuronal avalanches when excitation and inhibition are balanced.

We first show that *d2*, which was previously used to index distance-to-criticality in the firing rates of spiking neurons, also provides a reliable measure of distance-to-criticality in 2.5 s windows of EEG amplitude fluctuations (cf. Fig. S1). Progressively adjusting the control parameter by manipulating excitatory versus inhibitory connectivity (CE/CI), results in sharp transition from an absorbing, inactive phase (E/I < 1) to an autonomously active state (E/I > 1) (Gros, 2021; Fig. 2A) Moreover, the capacity for information transfer – an emergent property of critical systems (Fagerholm et al., 2016) – expands dramatically at this phase transition (Fig. 2B). Here, we measure mean mutual information in network output across time lags of τ = 1 to 25 timepoints for each CE-CI combination and plotted as a function of the corresponding network E/I balance derived from CE/CI. Importantly, *d2* (color-coded in Fig. 2A-B) faithfully indexes the distance of the control parameter to criticality. Mutual information peaks at the phase transition, where *d2* is the lowest, reflecting minimal distance-to-criticality.

**Figure 2.**
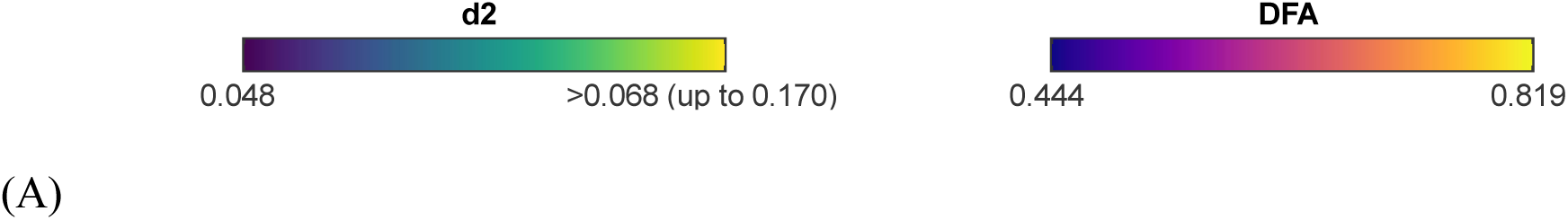
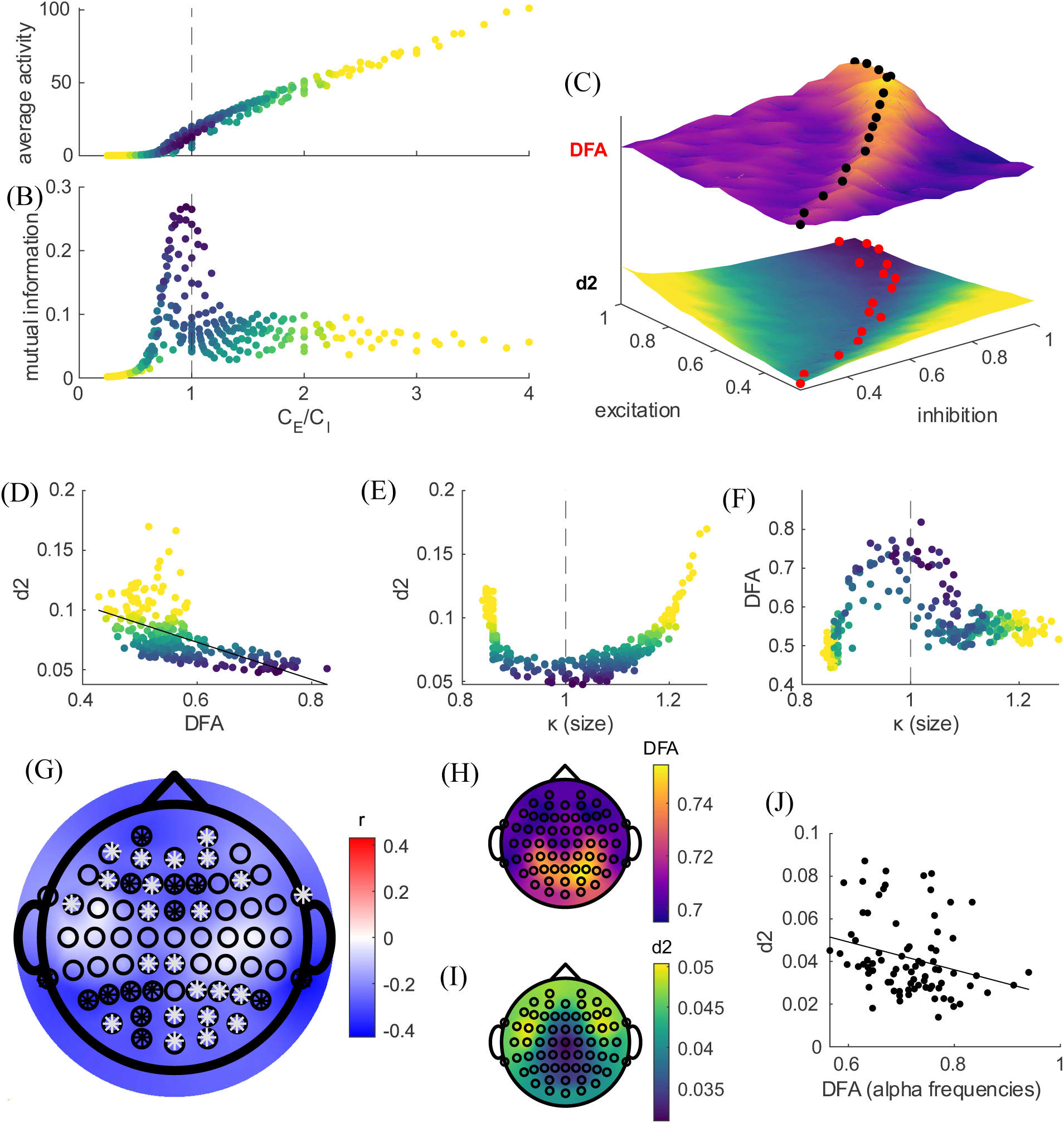
Benchmarking *d2* against ground truth simulations and resting-state data. The average network output signals from the CROS simulations (A) and the mutual information of the network output signals (B) as a function of time lag (τ =1 to 25) for each CE and CI combination, plotted with respect to CE/CI. Both plots are color-coded according to their corresponding *d2* values. The *d2* color bar is shown at the top of the figure, with the DFA color bar displayed on the right. (C) DFA (top) and *d2* (bottom) computed from the output signals across combinations of CE and CI, with values mapped according to the color bars shown above. Stacking the DFA grids on *d2* grids revealed overlapping region where *d2* was lowest (black dots on DFA grid) and where DFA was highest (<1; black dots on *d2* grid) across each CE condition. The overlap indicates that these conditions occurred when the CE and CI were balanced. (D - F) Scatter plots from the CROS simulations showing relationships between (D) *d2* and DFA, (E) *d2* and κ, and (F) alpha-band DFA and κ across all CE and CI combinations. (G) Pearson correlation between *d2* and DFA (averaged across alpha frequency band) in resting-state EEG, showing channels with higher correlations. White asterisks indicate *p* < 0.05; black asterisks indicate *p* < 0.05 after FDR correction. (H) Average resting-state DFA and (I) *d2* across participants, visualized as scalp topographies across EEG channels. (J) Participant-level scatter plot of average DFA (averaged across alpha frequency band) and *d2* in the resting state, with values averaged across trials and channels.

Next, we validated *d2* by comparing it to more well-established criticality indices. As predicted, we find that *d2* is lowest where emergent properties are maximized, including long-range temporal correlations, measured here by detrended fluctuation analysis: DFA (Peng et al., 1994; Poil et al., 2012), and scale-free avalanche distributions, measured here by a parameter which captures the relative distribution of avalanche sizes: κ (Shew et al., 2009). As predicted, we find that the smallest *d2* values closely align with the largest long-range temporal correlations (Fig. 2C). Fig. 2C shows the overlap of CE and CI combinations that have the lowest *d2* values and the highest α-band DFA across each row of % excitation connection. Fig. 2D shows that *d2* decreases with higher alpha-band DFA across all CE and CI combinations, yielding a negative correlation of r = -0.56, p = 2.49 × 10^-22^. Fig. 2E and F further demonstrate the networks with the lowest *d2* and highest α-band DFA coincide with κ = 1, indicating that network configurations closest to criticality also produce avalanche size distributions most consistent with theoretical power-law scaling.

### 3.2 Distance-to-criticality predicts long-range temporal correlations at rest

Next, we tested whether distance-to-criticality (*d2*; Fig. 2I) correlates with DFA (Fig 2H) in resting-state EEG. DFA was computed across 10 logarithmically spaced frequency bands spanning delta to gamma, and correlations were evaluated using DFA values averaged within the alpha band. Negative correlations were observed predominantly in posterior regions (parietal and occipital channels) as well as along the midline, indicating that stronger long-range temporal correlations (higher DFA) were associated with operating closer to criticality (lower *d2*) in these regions (Fig. 2G). Across all electrodes, however, individual difference correlations remain strong (Fig. 2J; r(78) = -0.27; p = 0.02), further confirming that *d2* correlates with DFA in EEG data as well as in our simulations.

### 3.3 Distance-to-Criticality and Pupil Dilation Track Task Demands

Distance-to-criticality (*d2*), calculated from timeseries of spiking neurons in rodents, varies with physical action (Sooter et al., 2025), motivating our hypothesis that distance-to-criticality tracks cognitive action as well. Pupil dilation has long been studied as an index of arousal, and effort expended during cognitive tasks, typically exhibiting a phasic increase during cognitive tasks that scales with demand (Jos et al., 2025; Kahneman, 1973; van der Wel & van Steenbergen, 2018). Here, we tested the prediction that pupil dilation correlates positively with *d2* during performance of a cognitive task.

We examined these predictions in the context of a task-switching paradigm, in which we replicate well-established findings that adapting to new task rules is more demanding than repeating the same task. As anticipated, compared to “Regular” trials where participants repeat the same task rules on roughly 5 out of 6 trials (e.g. odd-even judgments), they are less accurate on forced switch trials where they are required to switch (e.g., to high-low judgments; t(82) = 5.03, p = 2.81 × 10^-6^, p_FDR_ = 2.81 × 10^-6^), on voluntary switch trials in which they are presented with both tasks, and they can choose what they prefer to do (t(82) = 13.0, p = 1.21 × 10^-21^, p_FDR_ = 3.64 × 10^-21^), and on distractor trials in which they are presented with both tasks, but they are required to stick with the main task (t(82) = 7.07, p = 4.61 × 10^-10^, p_FDR_= 6.91 × 10^-10^; Fig. 3A). Reaction times mirror effects on accuracy, with participants responding faster in Regular trials relative to the non-Regular trials (forced-switch trials: t(82) = -22.9, p = 2.62 × 10^-37^, p_FDR_ = 2.62 × 10^-37^; distractor trials: t(82) = -27.2, p = -27.2 × 10^-43^, p_FDR_ = 7.49 × 10^-43^; voluntary-switch trials: t(82) = -37.0, p = 6.34 × 10^-53^, p_FDR_ = 6.34 × 10^-53^; Fig. 3B). Together, these behavioral outcomes suggest that non-Regular trials impose greater cognitive demand than more frequent Regular trials.

**Figure 3.**
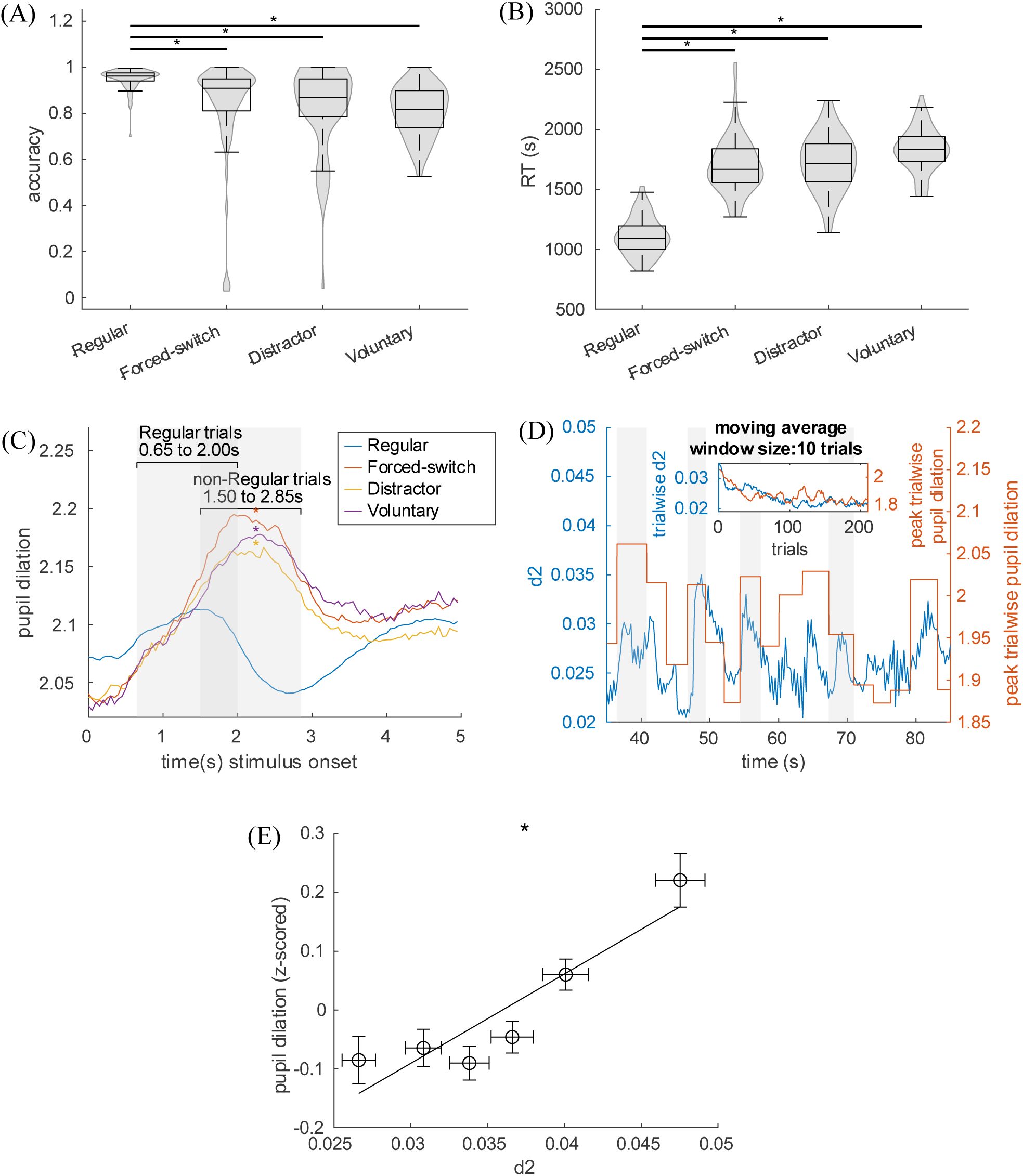
Deviation from criticality during pupil dilation. (A) Accuracy distribution and (B) reaction time (RT) distributions across trial types, with asterisk indicating significance after FDR correction. (C) Pupil diameter for different trial types: Regular, forced-switch, distractor, and voluntary-switch trials. Peak window for Regular trials (left grey region: 0.65 to 2 s post-stimulus) and non-Regular trials (right grey region: 1.5 to 2.85 s post-stimulus) were identified based on the peak of pupil response of the corresponding trial types. (D) The plot illustrates an example participant, showing *d2* (left y-axis) over time, computed using a 2.5-second sliding window with 90% overlap averaged across all EEG channels, alongside pupil diameter (right y-axis), which represents the mean peak-window pupil diameter for each trial type. Grey shaded regions indicate non-Regular trials. The inset displays post-stimulus *d2* values (0 to 2.5s after stimulus onset) together with the corresponding average pupil diameter in the peak window, smoothed using a moving average window of 10 trials. (E) Within-participant effects as illustrated by mean peak-window pupil diameter sorted by within-participant *d2* sextiles (all channels averaged), with mean values averaged across participants for each sextile. The error bar indicates the standard error of the mean across participants, while the asterisk indicates a significant effect of *d2* on pupil dilation from the mixed regression model.

Replicating prior work, we find that pupil dilation patterns track demand from trial to trial. We replicate well-established effects of trial-locked phasic pupil dilation (a task-evoked pupillary response, or TEPR; Fig. 3C) which peaks after participants respond on each trial (shaded gray regions mark the peak response window for Regular and non-Regular trials), and the peak is higher for more demanding trials. TEPR peaks are lower in Regular trials relative to forced-switch trials (t(47) = -7.95, p = 3.00 × 10^-10^, p_FDR_ = 9.00 × 10^-10^); distractor trials (t(47) = -7.04, p = 7.17 × 10^-9^, p_FDR_ = 1.08 × 10^-8^), and voluntary-switch trials (t(47) = -6.90, p = 1 × 10^-8^, p_FDR_ = 1.00 × 10^-8^; Fig. 3C).

Given that pupil dilation tracks task demands, the next question is whether *d2* exhibits corresponding variation alongside changes in pupil dilation. This prediction is motivated by prior work (Kardan et al., 2020) associating deviation from criticality with increased task demand, albeit at much slower timescales, across minutes of recording. Interestingly, the dynamics of *d2* and pupil dilation are remarkably similar across trials and trial types in two ways: they both track 1) trial-wise demand (Fig. 3D) and 2) trial number effects (inset of Fig. 3D). As illustrated for an example participant, *d2* fluctuations across trials corresponded closely with trial-by-trial peak-dilation patterns, with both measures increasing during non-Regular trials (gray shaded regions in Fig. 3D). Also, both pupil dilation and distance-to-criticality show a clear decrease across the task, declining as trials progressed, which may reflect practice effects or accommodation to task demands.

To test the relationship between *d2* and trial wise pupil dilation, we fit mixed-effects models regressing TEPR amplitude (*PD*) across all trials, onto post-stimulus *d2* (mean *d2* computing from sliding window estimates from stimulus onset to 2.5s post-stimulus), trial type (*k*), trial number (*t*), and their interaction as predictors, and included random intercepts and *d2* slopes across channels (*ch*) and participants (*P*):

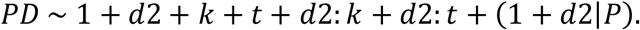

Effect estimates indicate that TEPR amplitudes increase reliably with *d2* (β = 0.0966, t(10015) = 5.67, p = 1.49 × 10^-8^), such that greater deviation from criticality corresponds to larger pupil dilation (Fig. 3E for within-participant effects). Pupil dilation also decreases as trials progress (inverse of trial number: β = -1.31 × 10^-3^, t(10015) = -21.6, p =1.49 × 10^-8^), and is larger during non-Regular trials compared to Regular trials (β = 0.120, t(10015) = 14.5, p = 7.01× 10^-47^). These pupil dynamics suggest not only that non-Regular trials are more demanding, but also that the beginning of a run of trials is more demanding and becomes less so as participants gain more practice. Importantly, the interaction of *d2* with trial number and trial type showed that the impact of deviation from criticality on pupil dilation was strongest in early trials (β = -4.31 × 10^-4^, t(10015) = -6.83, p = 9.23 × 10^-12^) and in non-Regular trials (β = 0.0270, t(10015) = 3.20, p = 1.38 × 10^-3^), highlighting how both practice and task demands may modulate the relationship between criticality and arousal.

### 3.4 Distance-to-criticality reflects task engagement and predicts inflexibility

We hypothesize that divergence from criticality reflects the degree of task engagement, or effort exerted during a task, beyond a passive response to task demands. Thus, we predict that effortful engagement progressively drives the brain away from criticality across a trial, as a function of load. To test the prediction, we calculated *d2* in a sliding window (2.5 seconds window, 98% overlap) across trials. We observe a clear pattern, that differentiates across trial types: during correct non-Regular trials, *d2* operates closest to criticality before trials start, then progressively diverges from criticality across the trial and peaks farthest from criticality after participants respond (Fig. 4A). Peak *d2* (0.5 to 1.0 s after stimulus onset) is higher than baseline (-1.0 to -0.5 s before stimulus onset: forced-switch trials: t(82) = 3.82, p = 3.00 × 10^-6^; distractor trials: t(82) = 2.12, p = 0.036; voluntary-switch trials: t(76) = 2.77, p = 7.00 × 10^-3^; voluntary-stay trials: t(75) = 3.68, p = 4.00 × 10^-4^), indicating that more demanding trials are associated with progressive deviation from criticality. In contrast, during Regular trials, scalp-averaged *d2* moves closer to criticality over the course of the trial. In fact, peak *d2* is lower than baseline, on average, (t(82) = -5.65, p = 9.00 × 10^-4^) on Regular trials. As a result, non-Regular trials end up further away from criticality than Regular trials during the post-stimulus peak window (t(82) = 4.90, p = 4.70 × 10^-6^).

**Figure 4.**
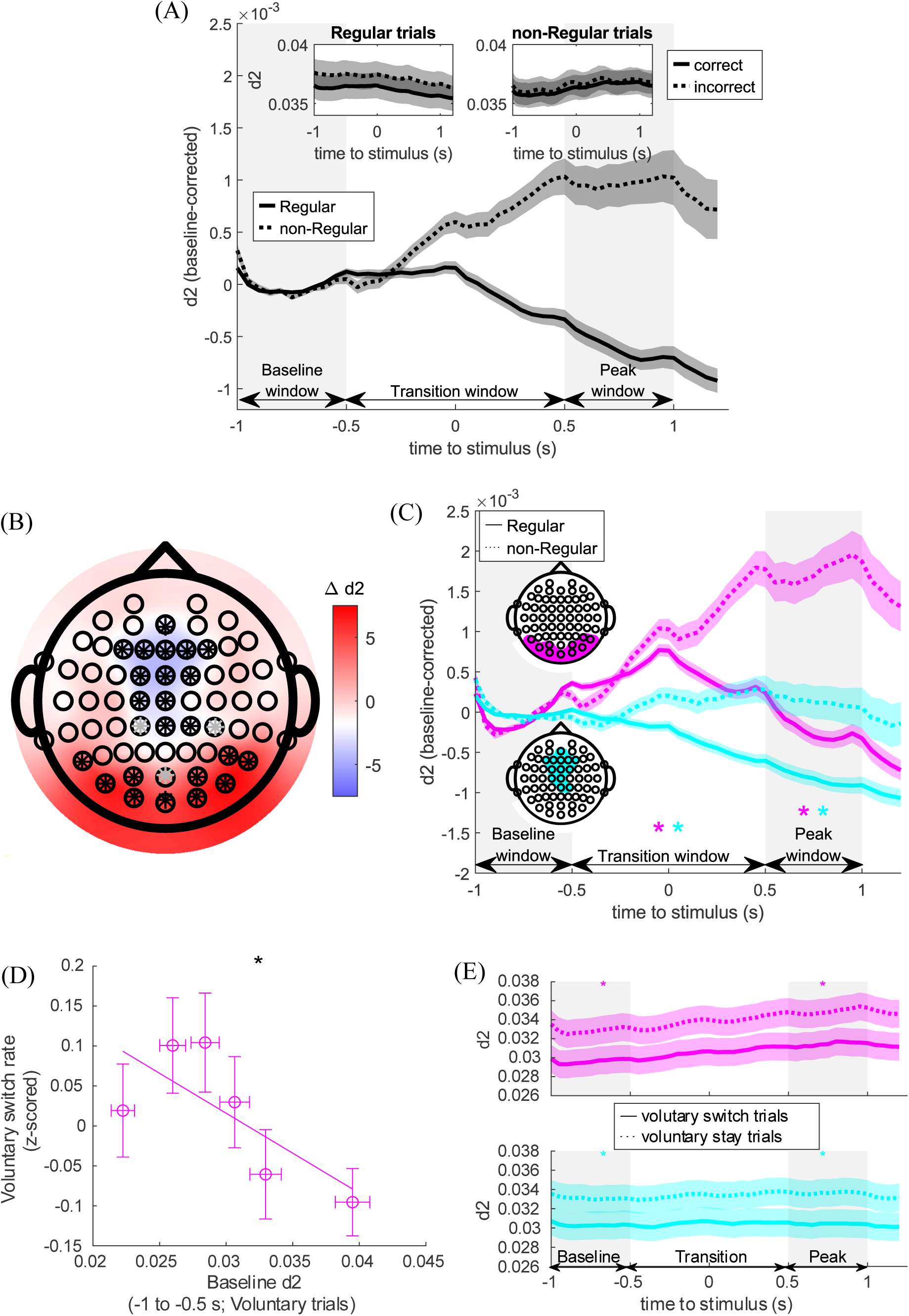
Higher *d2* predicts greater cognitive effort and diminished flexibility. (A) Sliding-window analysis of baseline-corrected *d2* (2.5s windows, 98% overlap) from -1.0s pre-stimulus to 1.2s post-stimulus onset, shown for Regular trials (solid black line) and non-Regular trials (dotted black line), averaged across all channels, correct trials, and participants. The inset displays the uncorrected *d2* for correct (solid black line) and incorrect (dotted black line) responses in Regular trials (top left) and non-Regular trials (top right). Shaded areas around the plot represent the standard error. (B) Topographic map of *d2* for Regular trials, comparing changes in *d2* from the baseline window to the transition window. Colors indicate t-statistics from a paired-test; white asterisk mark uncorrected p < 0.05, and black asterisk mark p_FDR_ < 0.05. (C) Same as panel A, but focusing on the sliding window of mean baseline-corrected *d2* across posterior (magenta) and midline (cyan) EEG regions separately. Asterisks indicate significant differences between Regular and non-Regular trials for each specific window and EEG region. (D) Mean posterior baseline *d2* and mean voluntary switch probability, each averaged within sextiles, where sextiles were defined by sorting trials based on *d2* values within participants, and grouping switch probability according to the exact ordering. Error bars represent the standard error. Asterisk indicates a significant effect of *d2* on voluntary switch rate from the mixed regression model. (E) Same as panel A, but plotted for *d2* only on Voluntary switch (solid line) and stay (dashed lines) trials for posterior (top plot, magenta) and midline (bottom plot, cyan) EEG regions. Asterisk on the top of each plot indicates significant difference between voluntary switch and stay trials for each specific window and EEG region. Shaded areas around the plot represent the standard error.

Interestingly, the effects of task engagement differ by region (Fig. 4B). Namely, comparing baseline (-1 to -.5 s pre-stimulus) and transition windows (-.5 to .5 s post-stimulus) on Regular trials reveals that posterior channels progressively deviate from criticality (transition *d2* > baseline *d2*; t(82) = 6.92, p = 6.09 × 10^-4^), while frontal midline channels get progressively closer to criticality (transition *d2* < baseline *d2*; t(82) = -4.53, p = 5.47 × 10^-4^; Fig. 4C). Considering the qualitatively distinct trajectories of these two sub-regions, selected from the remaining contiguous significant channels after FDR correction, subsequent analyses were conducted in each sub-region separately, with the cluster of significant midline EEG channels referred to as ‘midline’ (cyan, Fig. 4C) and significant parietal and occipital channels referred to as ‘posterior’ (magenta). During non-Regular trials, posterior regions (t(82) = 7.42, p = 9.61× 10^-11^) show progressive deviation from criticality during the transition versus the baseline window, but midline regions do not (t(82) = 0.95, p = 0.35). Also, at the peak window (.5 to 1 s post-stimulus), both posterior (t(82) = 3.94, p = 1.73 × 10^-4^) and midline regions (t(82) = 2.53, p = 0.0132) deviate further from criticality on non-Regular trials compared to Regular trials. Collectively, these results indicate that task engagement drives deviation from criticality as a function of task load, differentially across the scalp.

Criticality affords flexibility by virtue of emergent dynamical properties of critical systems, including heightened susceptibility (Beggs, 2022). Indeed critical dynamics predict greater cognitive flexibility in Go/NoGo (Simola et al., 2017) and perceptual bistability tasks (Pfeffer et al., 2018). As such, we predict that greater divergence from criticality is associated with diminished flexibility.

To test the relationship between criticality and flexibility, we tested whether individual differences in distance-to-criticality correlated with participants’ propensity to voluntarily task switch. This relationship was supported by a Wilcoxon rank-sum test, which indicates a significant difference in voluntary switch rates between participants in the lower versus higher baseline *d2* (median split), W = 1960, z = 1.77, *p* = 0.0390. In addition, a mixed-effects logistic regression model was fit to examine whether baseline *d2* activity predicted trial-wise switching on voluntary switch trials, with participant included as a random intercept. The model reveals a significant negative effect of baseline *d2* (β = -23.1, t(1513) = -2.09, p = 0.0360), suggesting that operating closer to criticality at baseline was associated with an increased probability of voluntary task-switching. Thus, we find that both individuals whose brains operate closer to criticality are more likely to voluntarily switch tasks, and that, within participants, voluntary task-switching is more likely when starting off a trial closer to criticality (Figure 4D - E).

### 3.5 Linking dynamic changes in criticality to within- trial performance

Emergent dynamics are proposed to maximize computational capacities by increasing susceptibility, information transfer, and storage capacity when operating near criticality (Gautam et al., 2015; Müller et al., 2025; Shew et al., 2011; Shriki & Yellin, 2016). Thus, a straightforward prediction is that operating closer to criticality makes participants faster and more accurate.

#### 3.5.1 Accuracy

To test this prediction, we regressed trial-by-trial accuracy (*A*) onto pre-trial, baseline criticality, using mixed-effects logistic regressions separately for midline and posterior EEG channels. Our model tested for the effects of baseline *d2* (*d2_b_*), trial number (*t*), trial type (*k*: Regular vs. non-Regular), and their interaction, while accounting for random intercept and random slopes of *d2_b_* and trial number across participants (*P*):

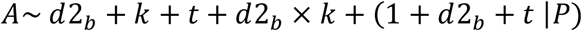

Regression results for midline EEG channels are listed in Supplemental Table S3, while those for posterior EEG channels are listed in Supplemental Table S4. Baseline midline *d2* is negatively related to accuracy (β = -0.161, t(21252)= -2.26, p = 0.0239, p_FDR_ = 0.0478; Fig. 5B), whereas baseline posterior *d2* is unrelated to accuracy (β = -0.0592, t(21485) = -0.897, p = 0.370, p_FDR_ = 0.370; Fig. 5A), suggesting that operating closer to criticality in midline channels, prior to a trial, improves performance. Trial type has a significant negative effect (e.g. in the midline channel analysis: β = -1.59, t(21252) = -29.0, p = 1.81×10^-181^, p_FDR_ = 1.81 ×10^-181^), indicating lower accuracy on non-Regular trials. The interaction between baseline *d2* and trial type was not significant in posterior channels (β = 0.105, t(21485) = 1.77, p = 0.0775, p_FDR_ = 0.0775) but is significant in midline channels (β = 0.190, t(21252) = 2.76, p = 5.35× 10^-3^, p_FDR_ = 0.0107), suggesting a region-specific modulation. In addition, accuracy significantly increases with trial number (e.g. in the midline region: β = 4.55×10^-3^, t(21252) = 8.66, p = 5.10 ×10^-18^, p_FDR_ = 5.10×10^-18^), indicating a general performance improvement with task progression.

**Figure 5.**
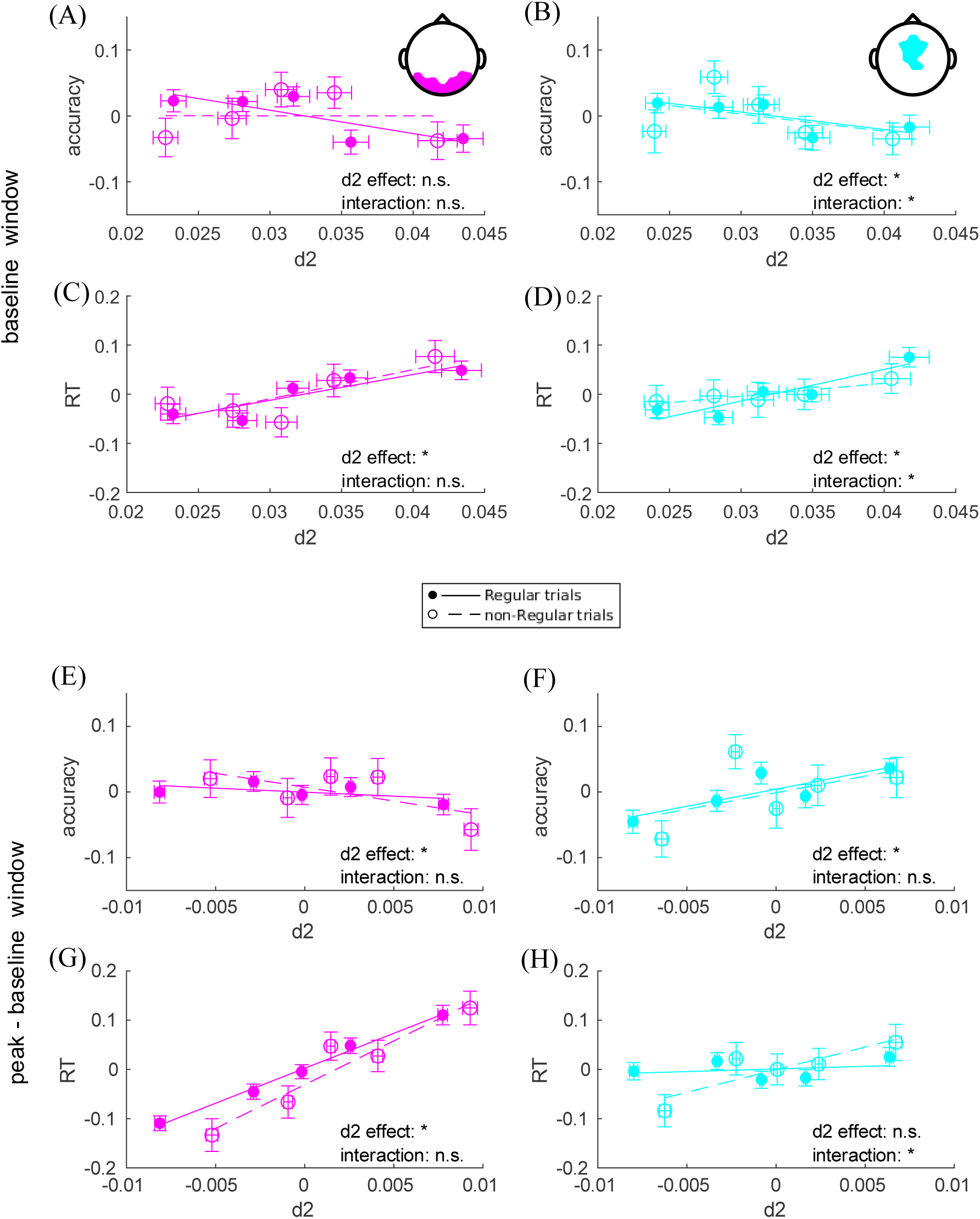
*d2* explains variability in reaction time and accuracy. Quintile plots of accuracy and RT sorted by within-participant *d2*, showing mean z-scored accuracy and RT and mean *d2* average across participants for each quintile. Panels (A) and (B) shows how accuracy corresponds to the posterior and midline regions during the baseline window, and panels (C) and (D) show how RT corresponds to the same regions during the same window. Panels (E) – (H) show the same analysis for accuracy and RT for the peak – baseline difference in *d2*, in the same order. Solid line indicates Regular trials, while dashed line indicates non-Regular trials. The pink line indicates the posterior region, while the blue line indicates the midline region. Asterisks indicate regression coefficient that are statistically significant for the corresponding predictor (Supplemental Tables S3 to S9). The error bar indicates the standard error of the mean across participants.

While baseline *d2* captures the brain state at the start of the trial, the difference between peak and baseline *d2* (*d2_diff_*) describes dynamic shifts away from criticality associated with task performance. Thus, we reasoned, *d2_diff_* might also be predict trial-wise accuracy. In separate mixed-effects models for midline and posterior channels, we regressed accuracy (*A*) onto trial number (*t*), peak-baseline *d2_diff_*, trial type (*k*), and their interaction (*d2_diff_* ×*k*) as fixed effects, and random intercepts and random slopes for *d2diff* and *t* by participant (*P*):

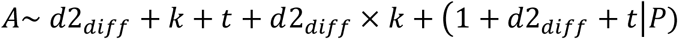

Regression results for midline EEG channels are listed in Supplemental Table S5, while those for posterior EEG channels are listed in Supplemental Table S6. Intriguingly, the direction of *d2_diff_* differed across regions: posterior *d2_diff_* is negatively associated with accuracy (β = -0.0928, t(21478) = -2.07, p = 0.0389, p_FDR_ = 0.0389; Fig. 5C), whereas midline *d2_diff_* is positively associated with accuracy (β = 0.0978, t(21245) = 2.12, p = 0.0337, p_FDR_ = 0.0389; Fig. 5D). The interaction between *d2_diff_* and trial type was not significant in either region (posterior: (β = 0.0464, t(21478) = 0.729, p = 0.466, p_FDR_ = 0.466; midline: β = -0.0535, t(21245) = -0.834, p = 0.404, p_FDR_ = 0.466). Taken together, the results suggest a functional dissociation between regions. During trial performance, a larger *d2_diff_* shift in midline channels, indicating transient increases in deviation from criticality, predict better accuracy. In contrast, larger *d2_diff_* shift in posterior channels predicts poorer task performance.

#### 3.5.2 RT

We also examined the effects of distance-to-criticality on reaction times (RT). First, we fit a mixed-effects models for baseline posterior and midline *d2* on RT (*RT*; on trials with correct responses only). In both models, we regressed RT onto trial number (*t*), baseline *d2* (*d2_b_*), trial type (*k*), preceding-trial accuracy (*A_t-1_*), and their interaction (*d2_b_* ×*k*) as fixed effects, and random intercepts and random slopes for *d2_b_* and *t* by participant (*P*):

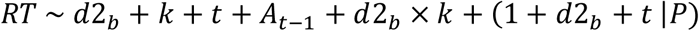

Regression results for midline EEG channels are listed in Supplemental Table S7, while those for posterior EEG channels are listed in Supplemental Table S8. Both posterior and midline models show that RT decreases with trial progression (e.g. midline model: β = -3.46×10^-4^, t(19496) = -2.60, p = 9.32 ×10^-3^, p_FDR_ = 0.0146), and increases following incorrect responses in the previous trial (midline: β = -0.318, t(19496) = -14.9, p = 2.98 × 10^-50^, p_FDR_ = 5.96 ×10^-50^). Trial type also has a significant positive effect in both regions (midline: β = 1.38, t(19496) = 101, p < 2.20 ×10^-16^, p_FDR_ < 2.20 ×10^-16^), indicating substantially longer RTs on non-Regular trials. Importantly, baseline *d2* positively predicts RT in both posterior (β = 0.0324, t(19705) = 2.51, p = 0.0119, p_FDR_ = 0.0119; Fig. 5E) and midline channels (β = 0.0598, t(19496) = 4.30, p = 1.73 × 10^-5^, p_FDR_ = 3.46×10^-5^; Fig. 5F), indicating that greater deviation from criticality is associated with slower responding. The interaction between baseline *d2* and trial type, on the other hand, reveals non-significant effects in the posterior region (β = 0.0233, t(19705) = 1.56, p = 0.119, p_FDR_= 0.119), and significant negative effects in the midline region (β = -0.0505, t(19496) = -2.95, p = 3.15 ×10^-3^, p_FDR_ = 6.30 ×10^-3^). The negative interaction at midline reflects the fact that the association between greater deviation from criticality and slower responses was attenuated under increased task demands in the non-Regular trials.

Similarly, we fit mixed-effects models regressing RT onto peak-baseline change (*d2_diff_*), in the posterior and midline regions, along with trial number (*t*), trial type (*k*), preceding-trial accuracy (*A_t-1_*), and the interaction with trial type (*d2_diff_* ×*k*) as fixed effects, and random intercepts and random slopes for *d2_diff_* and *t* by participant (*P*):

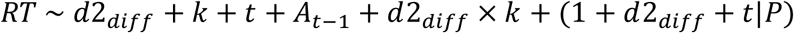

Full regression results for midline EEG channels are listed in Supplemental Table S9, while those for posterior EEG channels are listed in Supplemental Table S10. For posterior regions, greater peak-baseline *d2_diff_* deviation is associated with significantly slower responses (β = 0.0732, t(19698) = 7.66, p = 2.01×10^-14^, p_FDR_ = 4.02 ×10^-14^; Fig. 5G), whereas this effect is not significant in the midline region (β = 8.19×10^-4^, t(19489) = 0.0902, p = 0.928, p_FDR_ =0.928; Fig. 5F). The interaction between *d2_diff_* deviation and trial type is not significant in the posterior region (β = 0.0163, t(19698) = 1.00, p = 0.317, p_FDR_ = 0.317), but is significant in the midline region (β = 0.0451, t(19489) = 2.78, p = 5.41×10^-3^, p_FDR_ = 0.0108), indicating that at midline, the positive effect of peak-baseline *d2_diff_*on RT is amplified on non-Regular trials.

In sum, RTs decrease over trials, confirming typical learning and performance effects. For posterior channels, greater deviation from criticality, both at the baseline window and in the peak-baseline difference, is associated with slower responses, suggesting operating farther from criticality in posterior regions corresponds to reduced processing efficiency. For midline regions, higher baseline *d2* also predicts slower RTs, and the positive interaction between peak-baseline difference and trial type indicates that this slowing effect is amplified on non-Regular trials. This suggests that under higher task demand, greater task-evoked deviation from criticality in midline channels is associated with slower but more deliberate responding (as observed from the positive effect between peak-baseline *d2* and accuracy), consistent with increased engagement of effortful control.

### 3.6 Trait Expression of Criticality Across Rest and Task

While *d2* varies with task demands and task engagement, it also has trait-like properties, tracking individual differences across task states. We find that resting mean *d2* correlates positively with variance and mean of *d2* during task performance (Fig. 6). Taken together, these findings indicate that the degree of criticality expressed in the resting state (Fig 2G and 2J), reflects an individual trait that carries over into task engagement, such that the interindividual differences in resting-state dynamics predicts task-related deviations from criticality during active cognitive processing.

**Figure 6.**
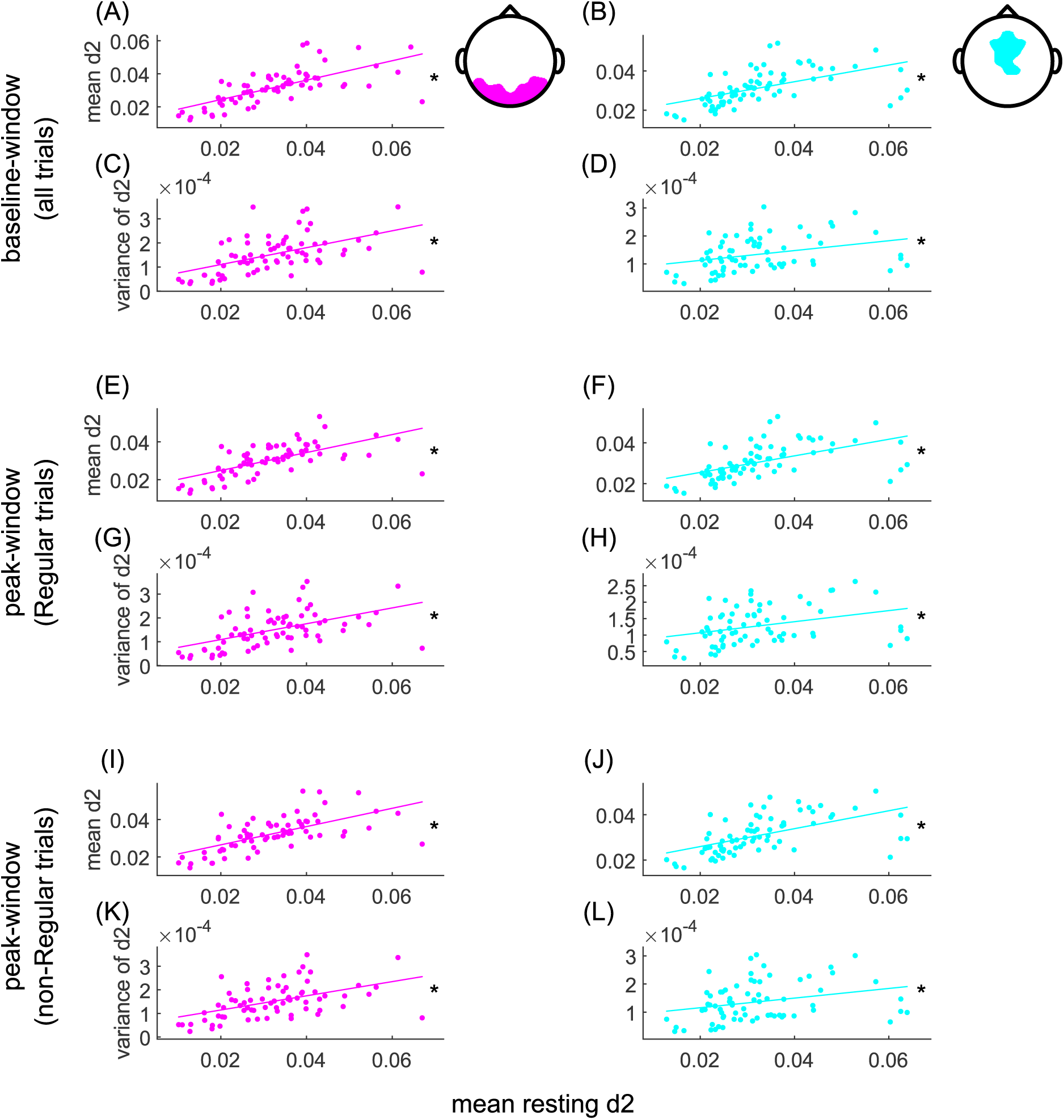
Resting *d2* correlates with task *d2*. Scatter plots show the relationship between resting-state d2 and on-task d2 in posterior (A, C, E, G, I, K) and midline (B, D, F, H, J, L) EEG regions. Mean and variance of *d2* during task performance are plotted against resting mean *d2* across on-task baseline and peak windows. Specifically, panels A – D depict mean and variance during the baseline window across all trials; panels E – H show mean and variance during the peak window in Regular trials; and panels I – L show mean and variance during the peak window in non-Regular trials. Asterisks indicate significant robust regression (p < .05; see Supplemental Table S11 for statistics).

## 4. Discussion

We demonstrate that brain criticality boosts cognitive flexibility, while cognitive load and cognitive effort drive deviation from criticality. To achieve this, we adapted the tRG-derived *d2* metric for application to EEG amplitude fluctuations. We begin by showing that *d2* faithfully tracks neural criticality at behaviorally-relevant timescales in both simulations and EEG recordings, providing a bridge between network dynamics and task-related behavior in capturing signatures of cognitive effort. Next, we examine the relationship between time-resolved *d2* and two complementary brain states supporting cognition with opposing predictions about criticality: task flexibility and task engagement, reflecting the ability to focus on, or switch between tasks as required. Both states have been linked with aspects of cognitive effort, yet they can be dissociable in their neural implementation (Jos et al., 2025; Riedel et al., 2022; Yan & Wang, 2025). Task flexibility may benefit from criticality in regions supporting information integration and adaptive evaluation of the environment, where slight perturbations can drive substantial state transitions (Gautam et al., 2015; Parr et al., 2024; Shriki & Yellin, 2016), allowing quick adjustment and adaptability to environmental changes. However, increased allocation of resources to a particular task drives networks away from criticality in regions mediating rule maintenance and focused attention (Fagerholm et al., 2015). *d2* provides a suitable index for capturing these states, as scale-free activity patterns emerge from networks operating near criticality, optimizing adaptability and flexibility, and deviations from this state reflect increasing reconfiguration costs associated with increased effort as the brain adapts to more demanding tasks (Shenhav et al., 2017). This perspective naturally connects to the traditional flexibility-stability tradeoff (Dreisbach et al., 2024; Dreisbach & Fröber, 2019; Egner, 2023; Hommel, 2015) in cognitive control, where moving away from brain criticality indicates a shift from a more flexible to a more stable and engaged brain state. Our findings support the hypothesis, revealing that distance-to-criticality (*d2*) correlates with task performance and participants’ switching behavior.

In simulations, we show that *d2* successfully tracks signal integration across timescales, and distance-to-criticality in terms of E/I balance (A.-E. Avramiea et al., 2025; Poil et al., 2012). Importantly, *d2* replicates traditional DFA and κ results across networks with varied E/I ratios. *d2* correlates negatively with DFA, showing that *d2* was lowest where DFA peaks. *d2* is also lowest where networks have κ values ≈ 1, at which point an avalanche size distribution is most consistent with power-law scaling (Poil et al., 2012; Shew et al., 2009). Complementing this, we showed that *d2* reached its lowest values during the network’s transition from a quiescent to an active phase. Such a network configuration also provides high mutual information indicating that information propagates across temporal scales when a network operates closest to criticality. Together, these results showed that *d2* parallels traditional metrics of criticality but, crucially, can be estimated over much shorter time windows (a few seconds), highlighting its suitability for examining trial-by-trial fluctuations. This is particularly important because the brain is not a stationary system (Galadí et al., 2021; Guan et al., 2020; Shen & Lin, 2019): neural dynamics shift rapidly across trials, so metrics that require long, stationary recordings will miss transient deviations (Arazi et al., 2017; Chen et al., 2024; Pedroni et al., 2017) related to arousal, engagement, and task demands.

Applying *d2* to EEG in the task-switching paradigm reveals that brain dynamics shift away from criticality in response to increased task demands, (e.g. on more demanding non-Regular versus less-demanding Regular trials). Regular trials occur far more frequently, providing a familiar context. Moreover, Regular trials require no rule changes thus requiring fewer cognitive operations, resulting in better performance. By contrast, non-Regular trials require participants to either switch task rules, which increases cognitive control demands (Bartek et al., 2023; Yeung et al., 2006), resist task-relevant distraction (Graydon & Eysenck, 1989; Lorenc et al., 2021; Papadopetraki et al., 2025), or exercise decision agency during voluntary-switch trials (Richardson et al., 2020), each of which imposes additional workload. Pupil dilation, a classic measure of cognitive effort (van der Wel & van Steenbergen, 2018), reinforces the interpretation that non-Regular trials are more demanding, and that *d2* was sensitive to demand. Trial-wise pupil dilation increases on non-Regular trials and positively correlates with *d2* values (i.e., increased distance-to-criticality) across all trials. Also consistent with the interpretation that *d2* tracks demand, we find that both *d2* and pupil dilation decrease across trials, perhaps because of accumulating practice effects, reducing effective demand across trials. Consistent with the inference that cognitive demand shifts E/I balance and thus distance-to-criticality, prior work showed that increased task demands (responding to less familiar tasks) induces a steeper 1/f slope in EEG, reflecting transient inhibitory engagement that interrupts ongoing processing, parallel to the effort-related deviation from criticality as observed in *d2* (Gyurkovics et al., 2022).

Beyond overall engagement-related shifts in *d2* during non-Regular trials, we observed a functional dissociation between midline and posterior regions in how distance-to-criticality evolves across Regular trials. Although both regions deviate from baseline criticality in non-Regular trials, their pattern diverges when task demands are lower. In posterior channels, *d2* briefly moves away from criticality after stimulus onset, before returning towards criticality within the same trial, consistent with a transient shift to support early sensory integration (Makeig et al., 2002). In contrast, midline *d2* shifts closer to criticality during Regular trials following stimulus presentation, suggesting that these regions approach a more efficient, low-effort state when task goals are well learned and stable (Haith & Krakauer, 2018). Together, these patterns indicate that within-trial deviations from criticality serve distinct computational functions across regions, highlighting complementary mechanisms for balancing efficiency and control during task performance. This finding challenges the idea of a single, uniform critical point for the entire brain and instead aligns with prior work showing that brain regions exhibit distinct and independent critical dynamics (Bansal et al., 2021; Liu et al., 2025). Bansal et al. (2021), for example, showed that avalanche dynamics are scale-invariant across rest and task but exhibit pronounced individual-and region- specific variability in brain criticality, supporting the view that brain dynamics operate within a critical-like regime composed of functionally dissociable regional processes rather than a single global critical state.

Regional dissociation in d2 is further supported by its relationship with task performance. During trials, posterior and midline regions progressively deviate from criticality, with opposing effects on trial accuracy and reaction times. In posterior areas, operating closer to criticality is associated with higher accuracy and faster responding, consistent with the idea that near-critical dynamics support efficient sensory integration. A similar result was shown in prior hippocampal work (Habibollahi et al., 2026), revealing that cognitive load shifts networks toward criticality, allowing maximal dynamical range for information transmission. Conversely, a greater within-trial shift away from criticality is associated with higher accuracy in midline regions, suggesting that control-related systems (Eisma et al., 2021) may require more stable, less susceptible dynamics to maintain task rules effectively. This opposite pattern indicates that criticality and performance are linked in a region-dependent rather than a global manner. Operating closer to criticality at baseline, prior to trial onset, predicts better performance in both regions. While this may seem at odds with a greater shift away from criticality predicting better performance in the midline during a trial, these findings reflect different phases of processing, with the latter shifts in the peak-window from its baseline reflecting how each region performs its functional role once active processing is engaged, across trial-by-trial variation in task demands.

We note that dissociable effects of posterior versus midline shifts away from criticality on performance has implications for cognitive effort. While shifting away from criticality as a function of load may reflect passive adaptation to cognitive demands in posterior channels, a shift away from criticality that enhances performance in the midline channels is more consistent with an interpretation of active engagement. The relationship between midline dynamics and performance likely reflects intentional recruitment of control resources to enhance task performance, which involves implicit cost-benefit decision-making about cognitive effort (Aben et al., 2020; Berry et al., 2017; Cavanagh & Frank, 2014; Holroyd & Yeung, 2012; Shenhav et al., 2017). Together, these findings show that *d2* captures region-specific, moment-to-moment fluctuations in criticality that differentially support preparatory processes and passive versus active processing of task demands. Posterior regions achieve optimal performance by remaining near criticality, while midline regions dynamically adjust their distance-to-criticality according to task demands, reflecting the stability-flexibility tradeoff central to cognitive control.

Regarding *d2*’s association with task flexibility, participants select less-frequent (non-Regular trials) over more-frequent (Regular trials) task, during voluntary-switch trials, when operating close to criticality before the trial starts. This pre-stimulus distance-to-criticality effect is consistent with prior evidence showing that voluntary task switching is supported by anticipatory, top-down control processes right before stimulus onset, in contrast to the post-stimulus, bottom-up mechanism dominating in forced task switching (Kang et al., 2014). This suggests that neural dynamics near criticality just before the trial support choosing alternative options and possibly promoting exploration and efficient task-set switching (Parr et al., 2024; Toghi et al., 2024). Task flexibility might improve when the brain has enough time to relax back to criticality prior to the subsequent trial, as post-trial relaxation dynamics can affect subsequent switching behavior. Consistent with accounts of residual switch costs, increasing preparation time reduces switch cost, likely reflecting lingering task-set inertia or retrieval of prior stimulus-response associations (Evans et al., 2015; Gladwin et al., 2006; Rangel et al., 2023; Ritz & Shenhav, 2024). From our study, incomplete relaxation away from the prior task state may manifest as persistent deviations from criticality that constrain voluntary task-switching. Future research should explore how the rate at which criticality is restored shapes adaptive choices.

Beyond moment-to-moment fluctuations, *d2* also captures stable inter-individual differences. Resting-state *d2* shows a positive correlation with on-task *d2* across trial types. This trait-like stability suggests that *d2* not only reflects transient task-evoked adjustments but also indicates intrinsic neural operating tendencies, where individuals carry systematic traits near criticality into task contexts. This trait-like aspect of criticality explains why, for example, long-range temporal correlations measured during rest can predict individual differences in performance during working memory task (Herzog et al., 2024). Importantly, implications for performance depend on task demands. In this prior study, individuals with stronger long-range temporal correlations during rest performed worse when tasks required stable maintenance of task representations, but they were unrelated to performance when participants had to flexibly update task representations. Thus, trait-like individual differences in distance-to-criticality may predict performance across tasks, as a function of distinct processing demands. Such consistencies position *d2* as a promising marker of cognitive behavior, showing how neural networks operate with respect to scale-invariant criticality in revealing variability in cognitive effort, balancing engagement and flexibility with respect to task demands, across both resting and task-evoked conditions. While these findings provide a strong foundation, several unresolved questions still need to be addressed in future research.

### 4.1 Limitations and future directions

A primary limitation of this study relates to spatial resolution. Although scalp EEG demonstrated significant regional differences between posterior and midline areas in this study, scalp signals inevitably combine multiple cortical sources. This complicates identifying which specific source or functions relate to the *d2* dynamics observed in EEG (Lai et al., 2018; Snyder et al., 2018). Future research using source-localized EEG, MEG, or multimodal approaches with structural connectivity could better pinpoint the cortical or functional systems that most significantly influence the *d2* criticality signature.

Secondly, while we use mean-corrected windowing to address non-stationarity in short segments of signal envelope to compute *d2*, *d2* estimates still depend on stationary assumptions even within relatively brief trial epochs. In Supplemental analysis (Supplemental Figure S1), we explore the ability of *d2* to resolve distance-to-criticality with varying degrees of stationarity and find a tradeoff between the reliability offered by estimates derived from longer duration segments and the sensitivity offered by estimates derived from shorter duration segments. Future research could explore non-stationarity within trials during the task and examine how it impacts the understanding of criticality adaptation during sustained tasks or dynamic shifts in internal states.

The *d2* measure is derived from an AR(25) model applied to data, down-sampled to 512 Hz. Because AR models operate over lagged samples, the resulting estimates are tied to the temporal resolution of the data, which may limit direct comparability across datasets acquired at different sampling rates.

Additionally, a conceptual limitation of the study involves the directionality in deviation from criticality. While *d2* measures how far a brain is from criticality, it does not indicate whether this deviation signifies movement toward more subcritical (more rigid or stabilized dynamics) or supercritical-like (more hypersensitive) regimes. This distinction is especially important in midline findings, where greater deviation correlates with better performance. However, it remains unclear whether improved control results from suppression of fluctuations (subcritical-like) or increased responsiveness (supercritical-like). To clarify this, it may be necessary to incorporate independent measures (e.g. excitation-inhibition balance) to look at local dynamics.

Lastly, a possible limitation of the study is the interpretability of *d2* deviations for cognitive effort, as they may be influenced by factors that scale with task complexity but are not solely attributable to cognitive effort. For instance, switching to non-Regular trials might produce greater arousal, evident in pupil dilation, which could modulate network gain and independently affect the temporal autocorrelation structure, shifting network dynamics away from criticality. However, such a state may not specifically indicate effort-related resource allocation. Moreover, task switching involves shifting attention and engaging working memory with relevant information (Hélie et al., 2023; Wang et al., 2022; Westbrook & Braver, 2016), and these transient cognitive changes could also cause similar deviations in multiscale temporal structure, regardless of effort. While *d2* effectively tracked behavioral responses to task demand and choice flexibility, sensitivity might instead reflect cognitive processes that covary with effort but are not exclusively driven by it. Furthermore, although the task inherently varies in cognitive demand, the present study did not directly quantify effort sensitivity through demand avoidance, limiting the extent to which *d2* can be interpreted as reflecting the subjective cost of effort. Future studies should incorporate experimental designs that orthogonalize cognitive effort from other variables, enabling clearer mechanistic insights into how observed deviation from criticality modulates cognitive effort.

