## Supplemental files for "From criticality to cognitive effort: scale-invariant EEG dynamics supporting cognitive flexibility are suppressed by effort"

### Supplementary Information

#### Window-dependent sensitivity of *d2* to state changes.

To evaluate *d2*'s ability to capture both stationary dynamics and non-stationary transitions between states, we conduct simulations using autoregressive (AR) model-based time series with known ground-truth distance from criticality measures through calculating *d2*. Three time series with chosen AR properties are generated with fixed ground-truth *d2* values of 0.026, 0.038, and 0.065, respectively. The AR formulation ensures that each state remained stationary over time when considered alone.

To introduce controlled transitions, the three AR-based states (time series chunks) are concatenated to create a continuous time series where state transitions occur at predefined rates. Rate of state change (switching timescale in Figure S1) varies across 0.5 to 30 seconds. After randomly picking the starting state, the alternating state is randomly chosen from the two remaining states (excluding the current one), ensuring systematic switching between distinct stationary AR states. Although each state segment may be short at faster switching rates, the AR structure remains valid up to a factor determined by segment length, allowing for meaningful estimation of *d2* within sliding windows. Sliding-window *d2* is computed using window sizes (moving window duration in Figure S1) ranging from 0.5 to 30 seconds. We calculated classification accuracy using a nearest-neighbor approach, whereby *d2* values computed from sliding windows are assigned to the closest ground-truth *d2* state. Near state boundaries, classification reflects the dominant contribution of the longer segment within each window.

Figures S1 summarizes classification accuracy as a function of window duration and switching timescale across varying noise conditions. In the noiseless condition (Figure S1A), accuracy increases sharply at the smallest window sizes across all switching timescale, then gradually declines as window duration increases. When state transitions are rapid, *d2* achieves peak classification accuracy using relatively short windows ($< 3$ s). In contrast, for slower switching dynamics, peak accuracy occurs at longer window durations before eventually decreasing as window size continues to increase. This result suggests a balance point and tradeoff between temporal resolution and the stability of *d2* estimation. Figures S1B and S1C show that this relationship is preserved when adding Gaussian noise to the time series ($\sigma = 0.1$ and 0.2, respectively), although overall peak accuracy decreases with increasing noise. Nevertheless, accuracy remains above chance (0.33) in most cases indicating that while accuracy declines as a function of the mismatch between analysis timescales and the switching time scale of the generative mechanisms, d2 retains a capacity to distinguish states.

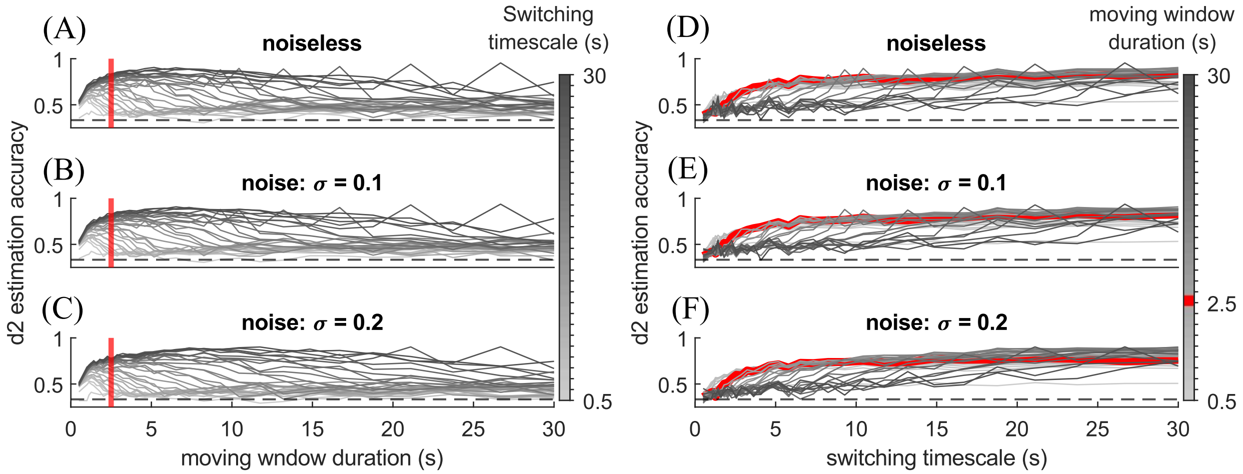


Figure S1. Classification accuracy of AR-generated time series with known distances from criticality (ground-truth *d2* for the three states: 0.026, 0.038, 0.065), based on nearest-neighbor matching of estimated *d2* values. Panels A-C plot accuracy as a function of window size for different rates of state switching, under (A) noiseless conditions, (B) moderate noise (σ = 0.1), and (C) higher noise (σ = 0.2). Panels D-F show the same results plotted as a function of state-switching rate for different window sizes, under the corresponding noise levels. The black dashed line represents chance-level performance (0.33), corresponding to one-third of the available choices.

Figures S1D-F show the additional analysis where classification accuracy is plotted as a function of switching timescales (x-axis), with separate curves for different window duration, matching the noise conditions in Figures S1A-C. Across noise levels, accuracy increases with slower switching rates when window sizes exceeded about 1 second, with smaller windows reaching their asymptotic peak accuracy sooner than larger ones. At a faster switching timescale, smaller window duration achieves higher accuracy than larger ones, emphasizing their benefit in tracking faster, non-stationary (slow) dynamics.

Together, these findings indicate that *d2* robustly captures both stationary dynamics and rapid sate transitions, with classification accuracy jointly influenced by window duration, switching timescale, and noise level. Based on these results, a window length of 2.5 s (red line in Figure S1) emerges as suitable choice for estimating *d2* during fast state changes, such as those occurring during force execution of trial presentation. This supports the use of short, second-scale windows for tracking criticality dynamics in neural signals that evolves over time.

#### Capturing network sensitivity to external inputs with *d2* and DFA

Aside from simulating the CROS model with 1% external spikes (*stimulus_size* = 25, approximately 1% of all neurons, or ~1.33% of excitatory neurons, were directly driven; Figure 2A-C), Figure S2 further show *d2* and DFA computed from CROS simulations across excitatory and inhibitory connection strength ($C_E$ and $C_I$) under 0% and 1.72% external spike input (*stimulus_size* = 0 and 43, respectively). AS shown in Figure S2A-B, the minimum of *d2* shifter toward the bottom-right direction of the $C_E$ and $C_I$ parameter space as external input increased, moving from regimes dominated by relatively greater excitation towards regimes with relatively stronger inhibition strength. This suggests that external drive increases network excitability, requiring stronger inhibitory coupling to restore balanced dynamics near criticality. In contrast, DFA showed a different sensitivity to external input. While the location of peak DFA remained relatively stable across CE and CI combinations, the overall magnitude of DFA decreased with increasing external spike input. This reduction likely reflects the introduction of externally driven activity that disrupts long-range temporal correlations by imposing additional short-lived variability. Together, these results highlight key distinctions between the two measures. *d2* sensitively captures how the locus of criticality shifts under changes in external drive, reflecting transient perturbations such as changes in trial events or sudden stimuli, whereas DFA primarily reflects a global reduction in long-range temporal correlation.

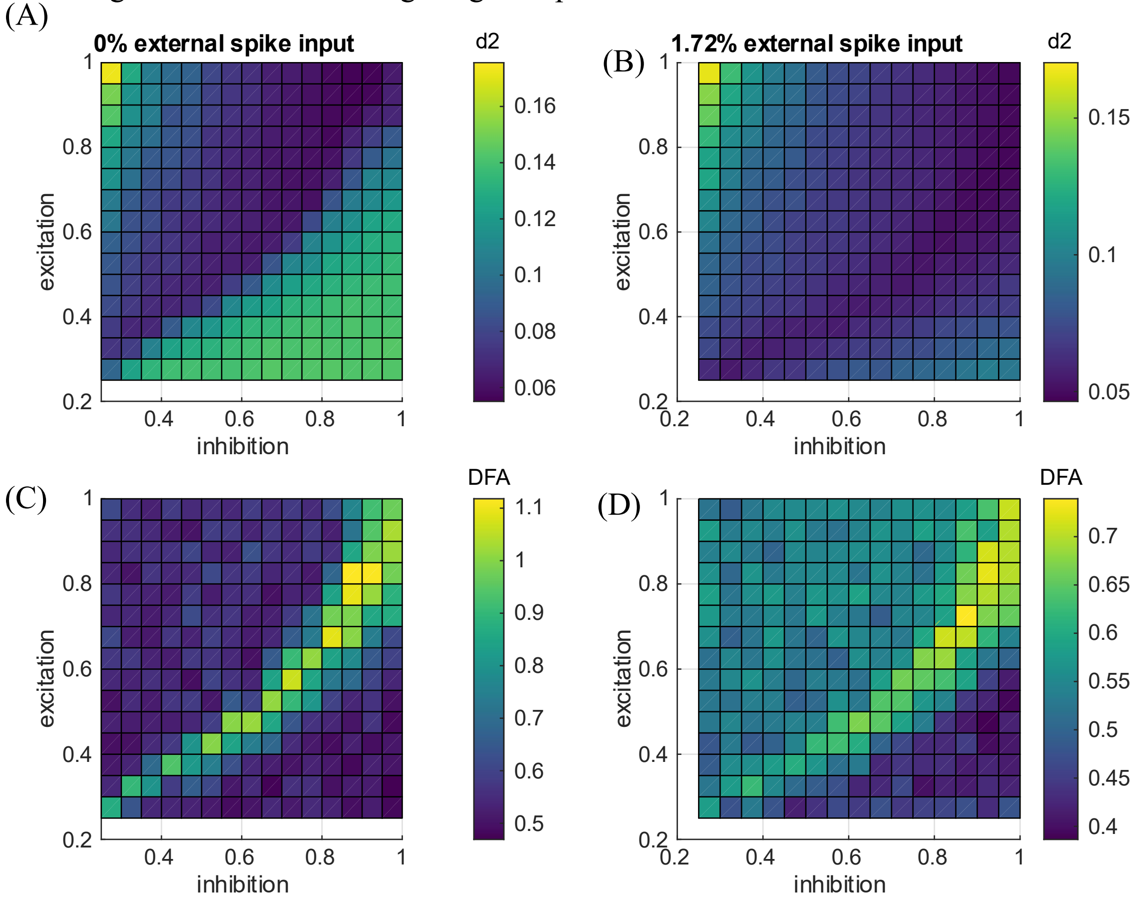


Figure S2. *d2* and DFA calculated for CrOS simulation across different combinations of $C_E$ and $C_I$ (A: *d2*; C: DFA) without external spike input and (B: *d2*; D: DFA) with 1.72% external spike input.

#### Regression results

This section reports the fixed-effects regression coefficients from the mixed-effects models described in the main manuscript. For ease of reference, Table S1 summarizes the notation used across all mixed-regression analyses in this section.

Table S1. Model notation and variable definitions.

| Notation | Definitions |
|---|---|
| $d2_b$ | Baseline *d2* |
| $d2_{diff}$ | Deviation of *d2* from baseline |
| *t* | Trial number |
| *k* | Trial type |
| $A_{t-1}$ | Preceding-trial accuracy |

Table S2. Fixed effect coefficients of the mixed-effect regression to examine the relationship between *d2* and trial-wise pupil dilation.

| Coefficient | β | SE | t-statistic | DF | p-value | Lower CI | Upper CI |
|---|---|---|---|---|---|---|---|
| Intercept | 0.145 | 0.133 | 1.09 | 10015 | 0.276 | -0.116 | 0.406 |
| *d2* | 0.0966 | 0.0170 | 5.67 | 10015 | $1.49\times10^{-8}$ | 0.0632 | 0.130 |
| *t* | $-1.31\times10^{-3}$ | $6.07\times10^{-5}$ | -21.6 | 10015 | $1.50\times10^{-101}$ | $-1.43\times10^{-3}$ | $-1.19\times10^{-3}$ |
| *k* | 0.120 | $8.30\times10^{-3}$ | 14.5 | 10015 | $7.01\times10^{-47}$ | 0.104 | 0.136 |
| *d2 : t* | $-4.31\times10^{-4}$ | $6.31\times10^{-5}$ | -6.83 | 10015 | $9.23\times10^{-12}$ | $-5.54\times10^{-4}$ | $-3.07\times10^{-4}$ |
| *d2 : k* | 0.0270 | $8.44\times10^{-3}$ | 3.20 | 10015 | $1.38\times10^{-3}$ | 0.0105 | 0.0435 |

Table S3. Fixed effect coefficients of the mixed-effect regression to examine the relationship between trial accuracy and pre-stimulus baseline *d2* ($d2_b$) in the midline channels.

| Coefficient | β | SE | t-statistic | DF | p-value | Lower CI | Upper CI |
|---|---|---|---|---|---|---|---|
| Intercept | 2.63 | 0.0980 | 26.9 | 21252 | $3.05\times10^{-156}$ | 2.44 | 2.82 |
| $d2_b$ | -0.161 | 0.0713 | -2.26 | 21252 | 0.0239 | -0.301 | -0.0213 |
| *t* | $4.55\times10^{-3}$ | $5.25\times10^{-4}$ | 8.66 | 21252 | $5.10\times10^{-18}$ | $3.52\times10^{-3}$ | $5.58\times10^{-3}$ |
| *k* | -1.59 | 0.0549 | -29.0 | 21252 | $1.81\times10^{-181}$ | -1.70 | -1.48 |
| $d2_b : k$ | 0.190 | 0.0682 | 2.76 | 21252 | $5.35\times10^{-3}$ | 0.0563 | 0.324 |

Table S4. Fixed effect coefficients of the mixed-effect regression to examine the relationship between trial accuracy and pre-stimulus baseline *d2* in the posterior channels.

| Coefficient | β | SE | t-statistic | DF | p-value | Lower CI | Upper CI |
|---|---|---|---|---|---|---|---|
| Intercept | 2.65 | 0.0956 | 27.7 | 21485 | $6.33\times10^{-166}$ | 2.46 | 2.84 |
| $d2_b$ | -0.0592 | 0.066 | -0.897 | 21485 | 0.370 | -0.189 | 0.0702 |
| *t* | $4.54\times10^{-3}$ | $5.07\times10^{-4}$ | 8.96 | 21485 | $3.56\times10^{-19}$ | $3.55\times10^{-3}$ | $5.53\times10^{-3}$ |
| *k* | -1.62 | 0.0544 | -29.8 | 21485 | $3.65\times10^{-191}$ | -1.727 | -1.51 |
| $d2_b : k$ | 0.105 | 0.0597 | 1.77 | 21485 | 0.0775 | -0.0116 | 0.222 |

Table S5. Fixed effect coefficients of the mixed-effect regression to examine the relationship between trial accuracy and post-stimulus deviation from baseline *d2* ($d2_{diff}$) in the midline channels.

| Coefficient | β | SE | t-statistic | DF | p-value | Lower CI | Upper CI |
|---|---|---|---|---|---|---|---|
| Intercept | 2.66 | 0.0961 | 27.7 | 21245 | $1.03 \times 10^{-165}$ | 2.47 | 2.85 |
| $d2_{diff}$ | 0.0978 | 0.0460 | 2.12 | 21245 | 0.0337 | $7.54 \times 10^{-3}$ | 0.188 |
| $t$ | $4.48 \times 10^{-3}$ | $5.13 \times 10^{-4}$ | 8.72 | 21245 | $2.88 \times 10^{-18}$ | $3.47 \times 10^{-3}$ | $5.48 \times 10^{-3}$ |
| $k$ | -1.61 | 0.0545 | -29.5 | 21245 | $8.09 \times 10^{-188}$ | -1.72 | -1.50 |
| $d2_{diff}: k$ | -0.0535 | 0.641 | -0.834 | 21245 | 0.404 | -0.179 | 0.0722 |

Table S6. Fixed effect coefficients of the mixed-effect regression to examine the relationship between trial accuracy and post-stimulus deviation from baseline *d2* in the posterior channels.

| Coefficient | β | SE | t-statistic | DF | p-value | Lower CI | Upper CI |
|---|---|---|---|---|---|---|---|
| Intercept | 2.64 | 0.0958 | 27.5 | 21478 | $3.28\times 10^{-164}$ | 2.45 | 2.83 |
| $d2_{diff}$ | -0.0928 | 0.0449 | -2.07 | 21478 | 0.0389 | -0.181 | $-4.72 \times 10^{-3}$ |
| $t$ | $4.65 \times 10^{-3}$ | $5.07 \times 10^{-4}$ | 9.16 | 21478 | $5.56 \times 10^{-20}$ | $3.65 \times 10^{-3}$ | $5.64 \times 10^{-3}$ |
| $k$ | -1.62 | 0.0548 | -29.5 | 21478 | $1.27\times 10^{-187}$ | -1.72 | -1.51 |
| $d2_{diff}: k$ | 0.0464 | 0.0637 | 0.729 | 21478 | 0.466 | -0.0784 | 0.171 |

Table S7. Fixed effect coefficients of the mixed-effect regression to examine the relationship between trial RT and pre-stimulus baseline *d2* in the midline channels.

| Coefficient | β | SE | t-statistic | DF | p-value | Lower CI | Upper CI |
|---|---|---|---|---|---|---|---|
| Intercept | 0.0496 | 0.0420 | 1.18 | 19496 | 0.237 | -0.0328 | 0.132 |
| $d2_b$ | 0.0598 | 0.0139 | 4.30 | 19496 | $1.73\times 10^{-5}$ | 0.0325 | 0.0871 |
| $t$ | $-3.46 \times 10^{-4}$ | $1.33 \times 10^{-4}$ | -2.60 | 19496 | $9.23 \times 10^{-3}$ | $-6.06\times 10^{-4}$ | $-8.53\times 10^{-5}$ |
| $A_{t-1}$ | -0.318 | 0.0212 | -14.9 | 19496 | $2.98 \times 10^{-50}$ | -0.359 | -0.276 |
| $k$ | 1.38 | 0.0137 | 101 | 19496 | $< 2.20\times 10^{-16}$ | 1.36 | 1.41 |
| $d2_b : k$ | -0.0505 | 0.0171 | -2.95 | 19496 | $3.15 \times 10^{-3}$ | -0.0840 | -0.0170 |

Table S8. Fixed effect coefficients of the mixed-effect regression to examine the relationship between trial RT and pre-stimulus baseline *d2* in the posterior channels.

| Coefficient | β | SE | t-statistic | DF | p-value | Lower CI | Upper CI |
|---|---|---|---|---|---|---|---|
| Intercept | 0.0425 | 0.0416 | 1.02 | 19705 | 0.306 | -0.0389 | 0.124 |
| $d2_b$ | 0.0324 | 0.0129 | 2.51 | 19705 | 0.0119 | $7.14\times 10^{-3}$ | 0.0577 |
| $t$ | $-3.24\times 10^{-4}$ | $1.33\times 10^{-4}$ | -2.44 | 19705 | 0.0147 | $-5.84\times 10^{-4}$ | $-6.36\times 10^{-5}$ |
| $A_{t-1}$ | -0.312 | 0.0211 | -14.76 | 19705 | $4.64 \times 10^{-49}$ | -0.353 | -0.271 |
| $k$ | 1.39 | 0.0136 | 102 | 19705 | $< 2.20\times 10^{-16}$ | 1.36 | 1.42 |
| $d2_b : k$ | 0.0233 | 0.0150 | 1.56 | 19705 | 0.119 | $-6.01\times 10^{-3}$ | 0.0527 |

Table S9. Fixed effect coefficients of the mixed-effect regression to examine the relationship between trial RT and post-stimulus deviation from baseline *d2* in the midline channels.

| Coefficient | β | SE | t-statistic | DF | p-value | Lower CI | Upper CI |
|---|---|---|---|---|---|---|---|
| Intercept | 0.0403 | 0.0416 | 0.970 | 19489 | 0.332 | -0.0412 | 0.122 |
| $d2_{diff}$ | $8.19\times 10^{-4}$ | $9.08\times 10^{-3}$ | 0.0902 | 19489 | 0.928 | -0.0170 | 0.0186 |
| $t$ | $-3.34\times 10^{-4}$ | $1.32\times 10^{-4}$ | -2.53 | 19489 | 0.0113 | $-5.92\times 10^{-4}$ | $-7.56\times 10^{-5}$ |
| $A_{t-1}$ | -0.319 | 0.0213 | -15.0 | 19489 | $1.17 \times 10^{-50}$ | -0.361 | -0.278 |
| $k$ | 1.38 | 0.0136 | 101 | 19489 | $< 2.20\times 10^{-16}$ | 1.35 | 1.41 |
| $d2_{diff}: k$ | 0.0451 | 0.0163 | 2.78 | 19489 | $5.41 \times 10^{-3}$ | 0.0133 | 0.0770 |

Table S10. Fixed effect coefficients of the mixed-effect regression to examine the relationship between trial RT and post-stimulus deviation from baseline *d2* in the posterior channels.

| Coefficient | β | SE | t-statistic | DF | p-value | Lower CI | Upper CI |
|---|---|---|---|---|---|---|---|
| Intercept | 0.0447 | 0.0415 | 1.08 | 19698 | 0.282 | -0.0367 | 0.126 |
| $d2_{diff}$ | 0.0732 | $9.57\times10^{-3}$ | 7.66 | 19698 | $2.01\times10^{-14}$ | 0.0545 | 0.0920 |
| $t$ | $-3.52\times10^{-4}$ | $1.30\times10^{-4}$ | -2.70 | 19698 | $6.89\times10^{-3}$ | $-6.07\times10^{-4}$ | $-9.66\times10^{-5}$ |
| $A_{t-1}$ | -0.312 | 0.0211 | -14.8 | 19698 | $2.05\times10^{-49}$ | -0.353 | -0.271 |
| $k$ | 1.36 | 0.0138 | 98.9 | 19698 | $< 2.20\times10^{-16}$ | 1.33 | 1.39 |
| $d2_{diff}: k$ | 0.0163 | 0.0162 | 1.00 | 19698 | 0.317 | -0.0156 | 0.0481 |

Table S11. Robust regression statistics for each mean d2 and variance d2 against resting mean d2 (Figure 6).

| EEG region | Trial type | Coefficient | β | SE | t-statistic | p-value |
|---|---|---|---|---|---|---|
| Posterior | All trials | Baseline mean d2 | 0.688 | 0.0628 | 11.0 | $7.76\times10^{-17}$ |
| | | Baseline d2 variance | $4.31\times10^{-3}$ | $6.00\times10^{-4}$ | 7.19 | $5.97\times10^{-10}$ |
| | Regular trials | Peak mean d2 | 0.596 | 0.0642 | 9.29 | $1.36\times10^{-13}$ |
| | | Peak d2 variance | $3.96\times10^{-3}$ | $6.17\times10^{-4}$ | 6.41 | $1.55\times10^{-8}$ |
| | non-Regular trials | Peak mean d2 | 0.582 | 0.0664 | 8.76 | $9.21\times10^{-13}$ |
| | | Peak d2 variance | $3.60\times10^{-3}$ | $6.18\times10^{-4}$ | 5.83 | $1.71\times10^{-7}$ |
| Frontal midline | All trials | Baseline mean d2 | 0.701 | 0.0556 | 12.6 | $9.11\times10^{-20}$ |
| | | Baseline d2 variance | $1.79\times10^{-3}$ | $6.47\times10^{-4}$ | 2.77 | $7.27\times10^{-3}$ |
| | Regular trials | Peak mean d2 | 0.667 | 0.0532 | 12.5 | $1.11\times10^{-19}$ |
| | | Peak d2 variance | $1.76\times10^{-3}$ | $6.11\times10^{-4}$ | 2.88 | $5.39\times10^{-3}$ |
| | non-Regular trials | Peak mean d2 | 0.678 | 0.0555 | 12.2 | $4.92\times10^{-19}$ |
| | | Peak d2 variance | $17.4\times10^{-3}$ | $7.28\times10^{-4}$ | 2.39 | 0.0195 |

##### 3.6.3 Recent Behavioral Outcomes Predict Changes in Neural Criticality

Beyond the primary results described in the main manuscript, we examine how participants' neural criticality is influenced by the most recent feedback. Because fluctuations in neural criticality may reflect adaptive updating in response to recent behavior, we conducted a regression analysis to test whether performance on the preceding trial shapes subsequent adjustments in *d2*. Mixed-effects regression is conducted separately on midline and posterior EEG channels to regress preceding-trial accuracy ($A_{t-1}$), trial number ($t$), and trial type ($k$: Regular vs. non-Regular trials) on the difference in *d2* between peak and baseline windows ($d2_{diff}$) on selected channels, including their interaction, while accounting for random intercept and random slopes of previous-trial accuracy and trial type across participants ($P$):

$$D2_{diff} \sim 1 + A_{t-1} + k + t + A_{t-1}{:}k + (1 + A_{t-1} + k|P).$$

The regression results are shown in Table S11. It shows that the preceding-trial accuracy significantly predicted the change in *d2* post-stimulus in the midline channels (β =-0.147, t(19490) = -5.09, p = $3.60\times10^{-7}$, $p_{FDRs} = 7.20\times10^{-7}$), but not in posterior channels (β = -0.263, t(19699)

= -1.07, p = 0.284, $p_{FDR}$ = 0.284). The results further show that post-stimulus *d2* in both midline and posterior regions deviates increasingly from baseline as the experiment progresses (midline: $\beta = 6.39 \times 10^{-4}$, t(19490) = 8.81, $p = 1.41 \times 10^{-18}$, $p_{FDR} = 2.82 \times 10^{-18}$; posterior: $\beta = 2.00 \times 10^{-4}$, t(19699) = 2.77, $p = 5.67 \times 10^{-3}$, $p_{FDR} = 5.70 \times 10^{-3}$), with posterior *d2* showing larger deviations during non-Regular trials ($\beta$ = 0.241, t(19699) = 2.99, $p = 2.79 \times 10^{-3}$, $p_{FDR} = 5.60 \times 10^{-3}$). This indicates that midline *d2* dynamics are influenced by prior trial performance, where an error in the previous trial shifts *d2* away from baseline criticality in the subsequent trial. Together, these findings suggest that criticality is not only modulated by ongoing task demands but is also affected by recent behavioral outcomes, particularly along midline regions, providing a mechanism through which past performance calibrates future neural engagement in the task.

Table S12. Fixed effect coefficients of the mixed-effect regression to examine the relationship between post-stimulus deviation from baseline *d2* in the midline channels and preceding-trial accuracy.

| **Coefficient** | **β** | **SE** | **t-statistic** | **DF** | **p-value** | **Lower CI** | **Upper CI** |
|---|---|---|---|---|---|---|---|
| Intercept | 0.0132 | 0.0269 | 0.489 | 19490 | 0.625 | -0.0396 | 0.0659 |
| $A_{t-1}$ | -0.147 | 0.0289 | -5.09 | 19490 | $3.60\times10^{-7}$ | -0.203 | -0.0903 |
| $t$ | $6.39 \times 10^{-4}$ | $7.26\times10^{-5}$ | 8.81 | 19490 | $1.41\times10^{-18}$ | $-4.97\times10^{-4}$ | $-7.82\times10^{-4}$ |
| $k$ | 0.118 | 0.0805 | 1.47 | 19490 | 0.142 | -0.0396 | 0.276 |
| $A_{t-1}$: $k$ | 0.0242 | 0.0789 | 0.307 | 19490 | 0.759 | -0.130 | 0.179 |

Table S13. Fixed effect coefficients of the mixed-effect regression to examine the relationship between post-stimulus deviation from baseline *d2* in the posterior channels and preceding-trial accuracy.

| **Coefficient** | **β** | **SE** | **t-statistic** | **DF** | **p-value** | **Lower CI** | **Upper CI** |
|---|---|---|---|---|---|---|---|
| Intercept | -0.0518 | 0.0254 | -2.04 | 19699 | 0.0410 | -0.102 | $-2.11\times10^{-3}$ |
| $A_{t-1}$ | -0.263 | 0.0246 | -1.07 | 19699 | 0.284 | -0.0744 | 0.0218 |
| $t$ | $2.00\times 10^{-4}$ | $7.22\times10^{-5}$ | 2.77 | 19699 | $5.67\times 10^{-3}$ | $5.82\times10^{-5}$ | $3.41\times10^{-4}$ |
| $k$ | 0.241 | 0.0805 | 2.99 | 19699 | $2.79\times10^{-3}$ | $4.69\times10^{-4}$ | $2.10\times10^{-3}$ |
| $A_{t-1}$: $k$ | $7.12\times 10^{-3}$ | 0.0785 | 0.0907 | 19699 | 0.928 | $-5.24\times10^{-4}$ | $1.04\times10^{-3}$ |
